# The complexity of micro-and nanoplastic research in the genus *Daphnia* – A systematic review of study variability and meta-analysis of immobilization rates

**DOI:** 10.1101/2023.03.24.534107

**Authors:** Julian Brehm, Sven Ritschar, Christian Laforsch, Magdalena M. Mair

**Author notes:** Contributed equally to this work.

## Abstract

In recent years, the number of publications on nano-and microplastic particles (NMPs) effects on freshwater organisms has increased rapidly. Freshwater crustaceans of the genus *Daphnia* are widely used in ecotoxicological research as model organisms for assessing the impact of NMPs. However, the diversity of experimental designs in these studies makes conclusions about the general impact of NMPs on *Daphnia* challenging. To approach this, we systematically reviewed the literature on NMP effects on *Daphnia* and summarized the diversity of test organisms, experimental conditions, NMP properties and measured endpoints to identify gaps in our knowledge of NMP effects on *Daphnia.* We use a meta-analysis on mortality and immobilization rates extracted from the compiled literature to illustrate how NMP properties and study parameters can impact outcomes in toxicity bioassays. In addition, we investigate the extent to which the available data can be used to predict the toxicity of untested NMPs based on the extracted parameters. Based on our results, we argue that focusing on a more diverse set of NMP properties combined with a more detailed characterization of the particles in future studies will help to fill current research gaps, improve predictive models and allow the identification of NMP properties linked to toxicity.

**Highlights:**

- Systematic review of NMP effects on the model system *Daphnia*
- Organismic, experimental and NMP properties influence observed effects
- *In silico* identification of traits likely linked to NMP toxicity (immobilization)
- More detailed standardized characterization of NMP needed to improve predictions

## 1. Introduction

Microplastic particles (MP, 1-1000 µm)[1] of different shapes and sizes have been found in almost all compartments of the environment, from ocean shores[2], open ocean waters[3], the deep sea[4] to terrestrial habitats[5] and freshwater ecosystems like streams[6,7] and lakes[7,8]. The same is likely for nanoplastic particles (NP, 1-1000 nm)[9], although their environmental detection is restricted due to analytical limitations. According to their origin, nano-and microplastic particles (hereafter NMPs) can be separated into two categories, primary and secondary plastic particles[10,11]. The first covers NMPs fabricated in these size ranges, like spherical micro-or nanobeads used for example, in commercial cosmetic products[12,13]. On the other hand, secondary NMPs are formed by the fragmentation of larger plastic debris, resulting in more complex feature combinations, including different sizes, shapes and surface areas[1,14,15].

NMPs are perceived as an emerging threat to the environment, and the number of studies investigating the occurrence and impact of NMPs has thus increased substantially during the last years[16]. Researchers use different model organisms from different habitats to study the toxicological effects of NMPs on biota. The planktonic crustacean *Daphnia magna*, part of the genus *Daphnia* (water flea), is among the main invertebrate model systems in aquatic environmental research. *Daphnia* spp. are pelagic filter feeders that play a key role in freshwater ecosystems by linking primary producers to higher trophic levels[17]. Consequently, species of the genus *Daphnia* are frequently used for basic research in freshwater ecology[18–21], as indicators for ecosystem health[22,23] and water quality[24], and the assessment of environmental risks of pollutants[25–28]. *Daphnia* are established test systems for both acute[29,30] and chronic[28,31] toxicity in the regulatory risk assessment of chemicals. As a standard test organism in ecotoxicology, the genus *Daphnia* has now also become a frequently used model for studying the impact of NMPs[29,30]. However, due to the complexity of *Daphnia* ecology, differing experimental conditions, and the diversity of NMPs[31,32], drawing general conclusions about NMP effects on *Daphnia* remains challenging. This becomes even more evident when considering that different *Daphnia* species respond differently to the same stressor[33,34], and there is even a high variability between *Daphnia* clones of the same species[35–37].

To draw conclusions on the impact of NMP on *Daphnia* spp., we need in-depth knowledge of the parameters that might influence the observed toxicities. These parameters range from the ecological/organismic level (e.g., tested species or clones) over the experimental level (e.g., temperature, type of food supply) to the NMP level (e.g., polymer type, shape, size, additives, surface charge, weathering).

Here, we review the literature on the effects of NMP on *Daphnia* spp.. We systematically screened studies for information on different study parameters (see mindmap in Fig. 1), including parameters of *Daphnia* ecology, applied experimental conditions, properties of the used NMP and measured endpoints. We illustrate the relationship between these parameters and the observed effects in a meta-analysis of mortality and immobilization rates extracted from the compiled literature. Using machine learning models, we investigated to what extent the compiled dataset can be used to predict the toxicity of untested NMP with different physico-chemical properties and identified parameters that were most important for producing accurate predictions. With this comprehensive approach, we identified gaps in the current research on NMP effects on *Daphnia* spp. and derived suggestions on which properties of NMPs might be responsible for the observed adverse effects. Based on our results, we give recommendations on how to improve future research on the impact of NMPs on *Daphnia*.

**Figure 1.**
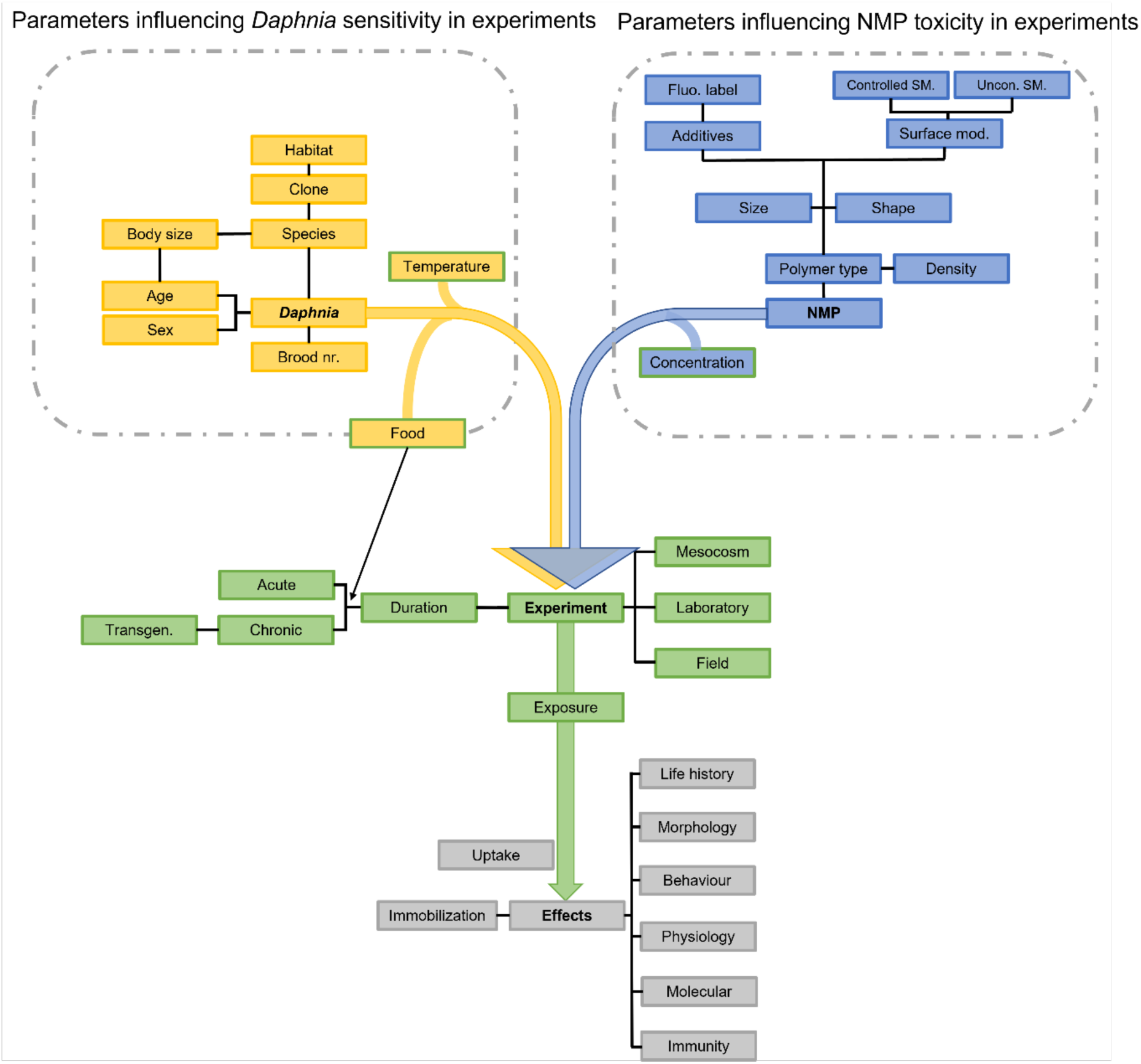
Mindmap on parameters influencing measured effects in experiments with *Daphnia* and NMP. The different levels are represented in color code: Parameters assigned to the organism and the organisms’ ecology are colored in yellow, parameters assigned to NMPs are colored in blue, parameters assigned to the experimental setup are colored in green, and the effects are colored in grey. In addition, parameters known to impact different levels are colored with an additional colored border according to the respective level. Abbreviations: Surface modifications (Surface mod.); Controlled surface modifications (Controlled SM.); Uncontrolled surface modifications (Uncon. SM.); Fluorescent label (Fluo. Label); Transgenerational (Transgen.).

## 2. Literature search and study parameters

We reviewed the existing literature on NMP effects on *Daphnia* spp. by conducting systematic searches on Web of Science (www.webofknowledge.com) using the following search terms: “*Daphnia* microplastic”, “*Daphnia* micro plastics”, “*Daphnia* nanoplastic” and “*Daphnia* nano plastics”. The last search was conducted on 18 February 2022 (239 hits). The titles and abstracts of all search results were screened to fulfil the following criteria: Only studies investigating NMP effects on *Daphnia* spp. were included; studies dealing with other micro-or nano particles were excluded; review articles, book chapters and non-English records were also excluded (101 remaining studies). For each article considered relevant, we extracted study parameters belonging to the four predefined main levels (visualized in the mindmap in Fig. 1): (1) *Daphnia* ecology/organismic level (yellow), (2) Experimental setup (green), (3) NMP characteristics (blue) and (4) Effects (grey).

### 2.1 Daphnia biology/organismic level

For the first main level, *Daphnia* biology/organismic level, information was collected on aspects of the test organisms that potentially impact outcomes in toxicity experiments. For each publication and experiment, we noted which species of *Daphnia* was used and if information about the used clone (yes/no), the habitat where it originates from (yes/no), sex (female/male), age (age of the test individuals), body size and brood number was given. We further noted food source (type of the food source) and temperature conditions (during the experiment and animal breeding).

### 2.2 Experimental setup

For the second main level, experimental setup, we noted whether the study/experiment was conducted under laboratory conditions, in mesocosm setups or in the field. Concerning the experimental time frame, we distinguished between acute toxicity tests (duration ≤96h), chronic toxicity tests (duration >96h) and transgenerational experiments where *Daphnia* were exposed to NMPs for more than one generation. In addition, when experiments with additional stressors (like added chemicals) were performed, only data on NMP exposure was extracted.

### 2.3 NMP characteristics

For the third main level, NMP characteristics, information regarding the plastics and, eventually, the non-plastic control particles used in the experiments were extracted. In detail, we examined information on the polymer type, density, size, shape (fibers, fragments, spheres), concentration (concentration is given yes/no, and measurement unit particles per volume or weight per volume), and if further physico-chemical properties are reported. These included NMP surface properties, additives melt-blended into the polymer and fluorescent labelling. For the physico-chemical properties, it was noted whether the information was given (e.g. information on surface modification yes/no) and which kind of properties were stated (e.g. surface modification carboxylated). We discriminated between uncontrolled surface modifications and controlled modifications. Uncontrolled surface modifications include modifications of the particles’ surface by milling, artificial weathering (using artificial UV light) or incubating the particles in biotic environments prior to use in toxicity tests[38,39]. In contrast to these “non-directional” and, therefore, uncontrolled modification types, we defined controlled modifications as modifications that alter the MNP surface structure and/or chemical composition in a more specific and controlled way. These modifications are frequently used to improve wettability, the adhesion to other materials, or the interaction with the biological environment[40] and include carboxylation and amination. In addition, we also included here the pretreatment with chemicals that can coat the polymer surface, e.g., Tween 20 or 80. Tween 20 and 80 facilitate the suspension of NMPs in aqueous media without shaking steps or sonication[41]. If a specific trait (e.g. a specific polymer type) was only used in one of the studies/experiments, it was listed as “other”. For the fluorescent labelling, it was noted whether particles were labeled or unlabeled and whether the dye was specified (yes/no).

### 2.4 Endpoints/Effects

In addition, information on the analyzed effects (measured endpoints) were extracted. Endpoints included in our review are: uptake (ingestion of NMPs); life-history traits such as time to maturity or the time to release the first offspring (neonates); reproduction (i.e. the number of offspring); morphology (changes in body size or growth rate of exposed *Daphnia*); behavioral effects (changes in feeding or swimming behavior); molecular effects (changes in protein abundance, alteration of the transcriptome, production of reactive oxygen species (ROS), or detoxifying enzyme expression); immune responses (e.g. count of immune cells), effects on the physiology (e.g. changes in the heart beat rate) and mortality/survival/immobilization rates. Data were recorded when any of the above endpoints were considered/mentioned.

### 2.5 Meta-analysis

As mortality and immobilization rates were reported most frequently in the screened literature and since only these endpoints are used in acute tests, we used these data for a more detailed meta-analysis of effect sizes (87 studies). For each of these publications and each treatment (i.e., each concentration level and tested NMP), we extracted the number of individuals that were dead/immobilized and alive/mobile for treatment and corresponding particle-free control, respectively. If several NMP treatments were compared to the same control treatment, the control treatment outcome was used repeatedly. If the number of dead/immobilized and alive/mobile individuals was not stated directly, we extracted the mean ± standard deviation (SD; or standard error (SE), or confidence interval) proportion of dead/immobile individuals. In addition, the number of replicates (i.e., test vessels), the number of individuals per replicate (i.e., the number of individuals in each test vessel) and the total number of tested individuals (i.e., replicates times individuals per replicate) were noted.

The number of dead/immobilized and alive/mobile individuals was then calculated based on the total number of individuals and the mean proportion of dead/immobile individuals, if necessary. If the same individuals were monitored over longer periods (e.g., checked every day in chronic toxicity tests), the immobilization rate for the latest measured of either 24, 48, 72, 96 hours, or 21 days was taken, and all other time points were neglected (e.g., day 21 was used instead of day 23, if both were reported). If measurements were not reported for either of these time points, data for the latest measured time point were used (e.g. day 63 if none of the time points mentioned above was reported). If not stated in the text or tables, immobilization/mortality rates were extracted from figures using the R package *metaDigitise*[42]. If the raw data were provided in the supplementary online material or provided by the authors, immobilization/mortality rates were calculated directly from the raw data instead. In addition, the source of each data point (e.g., from the text, extracted from figure x, calculated from raw data) was noted.

Where possible, concentration measures were transformed to mg per ml following recommendations by Thornton Hampton et al. (2022)[43]. Samples for which concentrations were reported as parts per million and samples for which a transformation into mg per ml was impossible were excluded from the analysis. Log Risk Ratios (RR) were calculated for each treatment-control pair as a measure of effect size using the R package *metafor* version 3.0.2[44]. Differences among different experimental parameters, organism and NMP properties were illustrated by grouped regression plots, including regression lines from meta-analytic mixed-effects models that included a random intercept for study ID[44,45].

#### 2.5.1 Predictive Modelling

We applied *in silico* machine learning methods to predict immobilisation log(RR) to the data extracted from the literature to investigate whether the available data can be used to predict toxic outcomes of untested NMPs. In addition, we used explainable machine learning methods to derive candidate parameters associated with toxic outcomes. To this end, we trained gradient-boosted regression trees to predict log(RR) based on NMP characteristics, experimental conditions and properties of the test organisms.

In total, 24 predictors were used for training, including 14 NMP characteristics, eight experimental conditions and two organism-related features. Boosted regression trees cannot handle NA values. To avoid biases resulting from filling up values by multiple imputation, additional factor levels were introduced for categorical predictors where reasonable. Samples not reporting the type of NMP modification were defined as “none”. For binary predictors (whether food was provided, particles were modified, UV weathered, fluorescent or coloured), NMP colour and NMP surface charge, NA values were defined as “not reported”. The remaining NA values were filled up by multiple imputation using *missRanger*[46,47]. Numeric predictors were scaled. One-hot-encoding finally resulted in 65 features for training.

Boosted regression trees were trained in R using *xgboost*[48]. A nested 5-fold cross-validation approach was applied for hyperparameter optimization, testing 50 different hyperparameter combinations selected randomly from a hyperparameter grid. The hyperparameter combination minimizing the residual mean squared error (RMSE) for predictions on test datasets was then chosen for the final model, which was trained on the complete dataset. To benchmark model performance, the same optimization procedure was repeated to establish a baseline model, i.e. a model trained on the same dataset but with log(RR) values shuffled randomly. The performance of the true data model was evaluated by (i) calculating the proportion of the variance explained by the final model and (ii) by comparing the predictive performance (by means of RMSEs from simple 10-fold CV using the respective optimized parameter sets) of the true data model to the baseline model. The final model was used to derive feature importances for the predictors (gain, cover and frequency).

All analyses were done in R version 4.2.2[45].

## 3. Results and discussion

In total, our web search found 101 articles (MP: 62; NP: 39) considered relevant for being included in our review (Fig. S1). The reviewed studies covered various experimental conditions, different test organisms and a diverse mix of NMP properties. 87 studies reported mortality/survival or immobilization effects and were considered for being included in the meta-analysis. Data extraction was finally possible for 59 of them.

For each parameter presented in Fig. 1, we will in the following summarize and discuss the impact it might have on experiments with *Daphnia*. In addition, we will discuss which NMP properties are possible candidates for eliciting adverse effects. There are fundamental differences between MP and NP, as the properties of MP cannot be easily extrapolated to NP particles and vice versa [49]. Furthermore, NP particles exhibit a nonlinear change in their properties compared to MP particles, requiring extensive re-evaluations of their behavior in the environment, formation mechanisms, and effects on organisms. To emphasize differences in parameter coverage between these two size classes, we thus report the results separately for MP and NP.

### 3.1 Daphnia biology/organismic level

In this section, we discuss the possible impacts of species characteristics such as body size, clone, and the chosen brood, as well as the impacts of different qualities/quantities of food supply and the temperature on the outcome of toxicity experiments with *Daphnia* using NMP.

#### 3.1.1 Body size

In all screened studies, body size is not determined for the organisms introduced into the experiment. However, body size of the daphnid determines, among other factors, the size range of particles that can be ingested (determined by the filtering area and the mesh size of the filter appendages) and the filtration rate. The filtration rate and filtering area increase with increasing body length[50–52]. This implements that smaller *Daphnia* species may ingest fewer NMP particles than larger species over time, as due to the lower filtration rate, smaller species need more time to filter the same amount of water than larger species. This species specificity is emphasized by a study of Brendelberger and Geller (1985)[52], who found a positive correlation between body size and the filtering area of the filtering apparatus for the two species, *D. magna* and *D. pulicaria*. These findings indicate that the larger the *Daphnia* species are, the more particulate matter is consumed and the broader the size range of particles they can ingest. Further, the mesh size of the filter apparatus can increase with increasing body size, as shown for *D. cucullata*[52]. For NMP experiments, this means that the size of the particles ingested most likely changes depending on the animal’s size. For example, for *D. parvula,* it was shown that algal filtration rates were higher and increased faster with increasing body size than the bacterial filtration rates did (26 to 33% of algal rates)[53]. That indicates that the efficiency in capturing ultrafine particles decreases with increasing body size, at least within a species[52,53]. This study used *Chlamydomonas reinhardtii* (∼10µm)[54] as the algal food source and bacterioplankton (<1µm) as the bacterial food source[53]. Due to the food sizes used in this experiment, one could theoretically equate the algae cell size with that of MP-particle size and the bacteria cell size with that of NP-particle size. Therefore, smaller species or younger and, thus, smaller animals may have higher NP uptake rates, while larger/older animals may have higher MP uptake rates. This holds only true if the particles are not aggregated. However, in most studies chemicals are used to keep the particles in suspension (see chapter 3.3.4.2.2). Consequently, depending on the particle size used in NMP studies, the choice of species and age is crucial since variations in body size might impact the outcome of a study.

In addition to the properties discussed so far, the capacity to resist toxicants is also a size-dependent. Generally, smaller *Daphnia* species or younger animals are more sensitive to dissolved stressors than larger species or older animals[55], which might also apply to particulate stressors[33].

As body size is usually not determined at the beginning of the study, it can only be estimated by considering the age of the introduced animals as well as the species used (see chapter 3.1.1.3). The dependencies of body size and particle size that can be ingested in connection with the feeding rate, the size of the filtering apparatus and size-dependent toxicity make it challenging to compare toxicity studies when different species or life stages are used. The effects found in one species or life stage could be over-or underestimated if one tries to transfer the results to another species or life stage implemented in a similar experimental setup. Hence, we recommend to report on morphological parameters of the used daphnids also at the beginning of each experiment.

##### 3.1.1.1 Age

The age of a daphnid is a major determining factor of body size. The age of individuals used in the screened studies ranged from embryos inside the brood chamber to 21-day-old adults.

However, the neonate stage (≤24h) was by far the most frequently used stage to start an experiment (62% overall; 55% MP; 74% NP) (for an overview of test organism age, see Fig. 2A).

**Figure 2.**
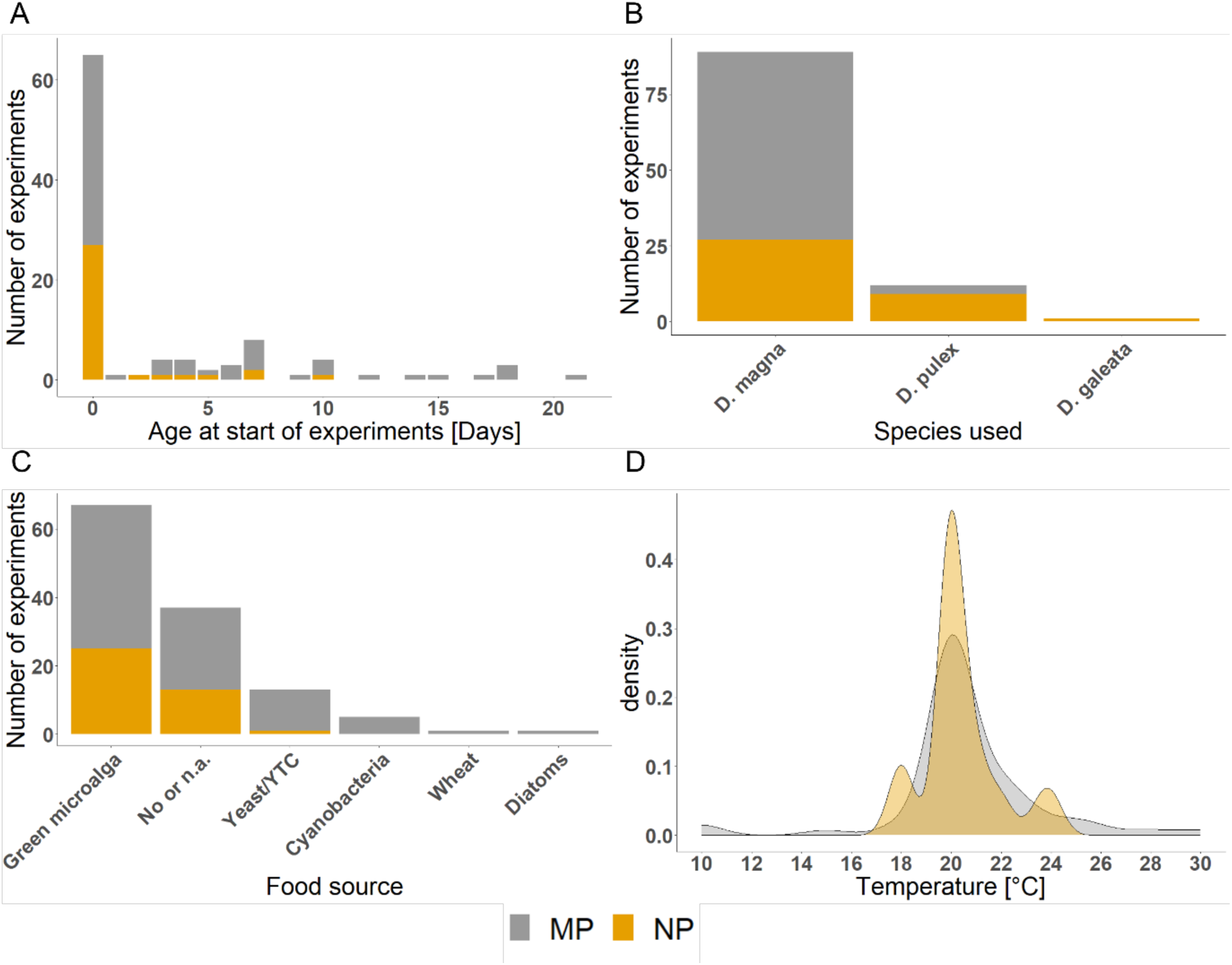
Traits linked to the *Daphnia* biology level. (A) Age of daphnids in days at the start of the experiment; (B) different *Daphnia* species used in experiments with NMP; (C) Different food sources used in experiments with NMP and *Daphnia*; (D) Density plot summarizing the temperatures used in experiments with NMP and *Daphnia*. Data are separated for studies dealing with MP (grey) and NP (orange). YTC: yeast, cerophyl and trout chow.

##### 3.1.1.2 Sex

All screened studies used females as test organisms, and none used males. Although males and females have the similar sizes as neonates[56], males grow slower than females[56] and reach smaller adult body sizes. Further, they have higher metabolic activity than females [56]. When using adult daphnids as test organisms, sex might thus be an additional factor influencing NMP toxicity[57].

###### 3.1.1.3 Species

In the studies we screened, 87% used *D. magna* (95% MP; 73% NP), followed by *D. pulex* (12% overall; 5% MP; 24% NP) and *D. galeata* (3% NP) (Fig. 2B; Fig. S2). Within *Daphnia*, body lengths between different species vary considerably. Adults range from mostly smaller than 2 mm in *D. galeata* to around 5 mm in *D. magna* (*D. pulex*: up to 3 mm)[58,59]. Further, the size ratio of adults to newborns varies significantly among cladocerans, whereby the size ratio of newborn to adult animals decreases with an increasing body size of the adult animals[60]. That means that larger *Daphnia* species produce smaller offspring in relation to their body length, which could negatively affect the comparability between acute (most often neonates are used) and chronic toxicity tests (endpoints measured on adults). As discussed in 3.1.1, these size differences between the species, the differences in offspring size, and the size ratio newborn/adult may have various implications on the test results in toxicity testing with NMP as body size has direct effects on, e.g., ingestible particle size.

*Daphnia* are filter feeders feeding on small, usually suspended particles[59]. Nonetheless, food uptake strategies can differ between species. While most *Daphnia* species feed exclusively on particles floating in the water column, *D. magna* can feed on organic substrate (algae, fine particulate matter) floating on or just below the water surface or settling through the water column (floating, surficial sediment, neuston)[61,62]. Further, they can stir up or feed on particles from the substrate (Fig. S3). Therefore, *D. magna* may encounter NMPs more often, at higher concentrations and likely also with different properties (e.g. larger positively buoyant particles[63]) than solely pelagic feeding *Daphnia* species. As a result, not only may the measured effects differ among *Daphnia* species due to differences in sensitivity but also because their different behavior results in different exposure doses.

###### 3.1.1.4 Clone

Given that daphnids reproduce parthenogenetic different clonal lines, consisting of different genotypes, are found in natural populations. Each clonal line shows different adaptations to its specific habitat. Therefore, responses of clonal lines differ in regard to biotic and abiotic stressors[64,65]. However, only 23% (32% MP; 8% NP) of the screened studies reported the used clone (here, parthenogenetic reproduction line), and the usage of a specific clone was restricted to specific working groups. Further, only in two studies clonal comparisons have been performed (both MP) [66,67].

The scarcity of annotated clones (genotypes) in the screened publications makes it difficult to compare the studies, as different clones of the same species may respond differently in ecotoxicity tests, even when used in the same experimental setup[36,66,68]. For example, Baird et al. (1991) analyzed the responses of five clones of *D. magna* to nine different chemicals in the form of a neonate mortality assessment (EC_50_) and found that responses can differ from one to two orders of magnitude[68]. There is also evidence of clone specificity in NMP research. A recent work by Sadler et al. (2019) found that the toxicity of MP exposure was clone specific and mainly related to an upregulation of hemocytes. Their results demonstrate how important it is to incorporate interclonal genetic variation into the assessment of NMP toxicity testing[66]. Therefore, sufficient information about the clonal line of a *Daphnia* species is necessary to understand the variability of effects analyzed for a species and the comparison between studies.

### 3.1.2 Brood number

We also evaluated annotations regarding the brood of *Daphnia* used in the experiments because offspring from different broods can differ in numbers, body size and fitness[69–71]. We found that only 20% of studies mentioned the brood they used, with the third brood being the predominant one (one MP study used the third to fifth brood in the experiments[72]).

Daphnids reproduce via cyclic parthenogenesis[73]. Given a life span (under perfect laboratory conditions) of around 150 days, as described for *D. magna*[74], and that an adult female may produce a clutch of eggs (brood) every three to four days[59], one daphnid may produce a vast number of broods during its life. In NMP research conducted with *Daphnia*, most studies use the third brood of age-synchronized animals for experiments since the number of offspring at this point is known to be rather consistent[70,71]. Also, in the OECD guideline 202, describing the acute toxicity assay for *D. magna* (testing for immobilization/mortality), it is specified that the animals should not be first brood progeny, to reduce the variability in, e.g. size classes between individual offspring and hence make the experiment more robust[75].

Besides the directive to only use animals from a specific brood onwards, the acute toxicity guideline (OECD guideline 202[30]) additionally states that animals should only be used when the culture is healthy, and no males and no resting eggs (ephippia) are produced. The last two factors, the production of males and resting eggs, need to be considered in an experiment since *Daphnia* change their reproduction mode in response to unstable environments and when environmental conditions deteriorate. Hence, when *Daphnia* switch to sexual reproduction, it implies that the culture used might be stressed or unhealthy[76–79]. To foster comparability between experiments, care needs to be taken when rearing the organisms for an experiment and the third brood of healthy offspring should be chosen for NMP experiments.

In addition to the test individuals’ characteristics, other abiotic and biotic factors in the environment may directly influence the impact of NMP on *Daphnia* in experiments. Critical are, for instance, food quality and quantity, and temperature.

### 3.1.3 Food supply

Daphnids can feed on various food sources like diatoms, green algae, cyanobacteria or organic debris (detritus)[80]. Most often, high food quantities according to OECD guideline 211[28] of differing quality were used in the screened experiments considered in this review (except acute experiments). In the studies we screened, food was provided in 54% of experiments (49% MP, 64% NP) and many different food sources were used (see Fig. 2C), with most of them being different species of green microalga (*Acutodesmus* sp., *Chlorella* sp., *Desmodesmus* sp., *Raphidocelis* sp.). In 30% of experiments, no food was supplied (acute toxicity tests) or the food source was not specified (33% MP; 28% NP). Besides algae, other food sources and supplements like yeast & YCT (yeast, cerophyl and trout chow) (10% overall; 14% MP, 3% NP), wheat (one MP study), and cyanobacteria (4% overall; 6% MP, 0% NP), were provided as food.

Differing food sources or limited food levels may affect the outcome of ecotoxicity studies, as starved individuals have fewer resources available for physiological defenses against stressors[81]. Several ecotoxicity studies demonstrated that limited food levels (in quality or quantity[75]) could increase the toxicity of stressors, i.e., pesticides[82], cadmium[83] or silver nanoparticles[84]. Poor food quality can be characterized by an insufficiency of a subset of essential resources, and poor food quantity can be characterized by an insufficiency of available energy and essential resources[85]. Concerning *Daphnia*, the availability of essential polyunsaturated fatty acids and phosphorus may determine the food quality[86,87], even between different green algae species[88]. For instance, Conine & Frost (2017) found lower toxicity of silver nanoparticles on *Daphnia* when feeding phosphorous rich compared to phosphorous-poor food[89]. Hence, to obtain high survival rates in control treatments, a high food supply of sufficient quality is usually provided in chronic laboratory ecotoxicity tests[90,91]. According to the OECD guideline 211, ratio levels between 0.1 and 0.2 mg/C/daphnid/day are sufficient, and the diet should preferably be living algal cells of, e.g., *Chlorella* sp.[28].

However, concerns have been raised about whether high food levels are optimal when aiming to assess risks under natural conditions. Stevenson et al. (2017) argue that *Daphnia* experience fluctuations in algal food supply in natural environments and that in standardized chronic toxicity testing, up to 400 times more food is present [92]. Aljaibachi & Callaghan (2018) showed that the presence of algae decreased the uptake of MP by *D. magna,* which was not in proportion to the relative availability[93]. Porter et al. (1982) observed that with increased food supply and overcollection of food particles, *Daphnia* responded with an increased rejection of food particles[94]. Geller & Müller (1981) found that if food quantities were high, *Daphnia* spp. switched to selective filter-feeding while at lower food levels, this selectivity was absent. In terms of experiments with NMP, this could mean that when little food is available in experiments, *Daphnia* may ingest all particulate material they encounter, while at high food availability, they may select what they ingest. For NMP experiments, this could mean that *Daphnia* are more likely to ingest NMP at low food concentrations than at higher ones. Considering annual fluctuations in *Daphnia* abundance as well as food availability, this might lead to discrepancies in NMP effects in natural environments compared to effects found in laboratory experiments[95]. High food levels in NMP studies may lead to an underestimation of effects on *Daphnia* in their natural habitat since daphnids probably consume less food in the environment as offered in experimental setups. Although standardized test protocols with high food supply are critical for comparing the effects of different pollutants in a standardized way, we believe that additional experiments with more limited, realistic food availability could be beneficial for gaining a better mechanistic understanding of processes found in contaminated environments.

### 3.1.4 Temperature

Standard toxicity tests are performed at a constant optimal temperature between 18°C and 22°C[27,28] (in practice, usually 20°C)[96]. In the literature we screened, most publications followed these recommendations, with most (42 MP; 26 NP) experiments conducted at temperatures between 18 – 22°C. Some studies, however, were conducted at more extreme temperatures going down to 10°C and up to 30°C (Fig. 2 D). For 12 MP and 10 NP experiments in the screened literature, no information on experimental temperature was provided.

Temperature is an essential experimental parameter, as it directly affects physiological processes, development and the daphnids’ behavior (e.g., swimming behavior)[97,98]. For example, Mc Kee & Ebert (1996) found that the body length of daphnids decreases with increasing temperature[98], and Nikitin et al. (2019) observed that *D. magna* increase their swim speed with increasing temperatures[99]. Changes in swimming behavior might then influence the encounter rate of *Daphnia* with NMP particles and thus change observed outcomes.

Further, temperature can directly influence the effects of toxicants. For example, there is a general trend that with increasing temperature, the toxicity of chemical compounds on *Daphnia* increases[96,100,101]. Heugens et al. (2003) state that an enhanced sensitivity at higher temperatures was the main driver for temperature-dependent toxicity. They add that a temperature correction may be necessary when translating toxicity data from the laboratory to the field[96]. That also accounts for NMP studies, where the toxicity of NMP has been shown to increase with increasing temperatures[102].

Overall, to gain better comparability between studies, we would recommend sticking to a temperature of ∼ 20°C in *Daphnia* experiments. However, additional experiments with higher or lower temperatures (or even temperature fluctuations) could be beneficial to gain a better mechanistic understanding of processes found in contaminated environments.

### 3.2 Experimental setup

#### 3.2.1 Study scope - Laboratory, mesocosm and field

All screened studies, except for one mesocosm study, were laboratory studies. Studies conducted under field conditions were not found.

Laboratory studies are easy to conduct and provide standardized data comparable among pollutants in a relatively short period of time. As a result, laboratory experiments serve as the basis for lower-tier regulatory risk assessment of chemicals.

Apart from the well-established laboratory studies, experiments simulating more natural systems are crucial to investigate impacts under more realistic conditions. A step towards this endeavor is the implementation of mesocosm experiments. Mesocosms simulate lentic aquatic habitats[103] and enable researchers to examine impacts on simplified aquatic communities. In the only mesocosm study we found, various parameters like the effects of MP on the species abundance of *Daphnia* and mayfly larvae (*Leptophlebia* spp.) and the distribution of MP in the water and sediment were analyzed[104], thus, combining hazard assessment with the assessment of exposure and particle fate. In addition, mesocosm studies enable the analysis of population-level effects and the impacts of pollutants on species interactions.

Conducting field experiments with NMP is not trivial; to our knowledge, no such attempts have been made so far. Nonetheless, it would be intriguing to look into the effects of NMP on *Daphnia* in a natural setting. Temperature, photoperiod, and nutrition are very variable under field conditions and may impact how NMP affects *Daphnia* populations[105]. Using field cages for daphnids seems promising for such a field experiment[105]. In their study, O’Connor et al. (2021) found that cages did not impede food flow and that the performance of clones in cages was equivalent to that in jars. With such an approach, it would be possible to study differences in life histories and population growth among lakes containing different amounts of NMP.

Although laboratory studies are essential for getting an overview on possible effects, we think mesocosm studies and field studies would be helpful and essential to enable the transfer of results from the laboratory to the possible impact of NMP in nature.

#### 3.2.2 Standard test procedures

##### 3.2.2.1 Duration

About two-thirds of experiments in the screened literature followed an acute toxicity test design (:< 96 hours of exposure; 61% overall; 58% MP; 67% NP), and about one-third used a chronic design (> 96 hours of exposure; 39% overall; 42% MP; 33% NP; with 3 MP and 3 NP studies using transgenerational approaches).

Acute toxicity tests with *Daphnia* are one of the most used bioassays for toxicity screening [106]. Here no food is added during the experiment. Standard endpoints are survival/mortality (LC50) and immobility (EC50)[27]. Advantages of acute toxicity tests are, for example, the rapid achievement of results, the possibility of high throughput methods, the simple experimental setup and good standardizability. Disadvantages include the limited number of endpoints and that sublethal effects are not covered.

Chronic toxicity tests with *Daphnia* usually last 21 days (as specified by the OECD), and standard endpoints are parental mortality and the number of offspring[28]. In chronic toxicity testing, daphnids are fed daily, and a complete water/particle exchange is performed every second day. An advantage of these tests is, for example, the ability to detect sublethal effects on many different endpoints and thus also to detect effects of non-acutely toxic substances. The main disadvantage is certainly the immense amount of work compared to the acute tests. A special case of chronic testing is transgenerational testing, where the effects of the same test substance are screened over multiple generations. This gives researchers the possibility to detect, for example, maternal effects, as well as adaptation effects.

In NP research, almost all screened studies stuck to the proposed exposure duration of 48h for acute and 21 days for chronic tests (see two distinct peaks in Fig. 3). In MP research, however, modifications of exposure durations occurred more often (see flatter and broader curve for MP in Fig. 3). Here, no clear distinction between acute and chronic experiments based on the duration of an experiment was possible.

**Figure 3.**
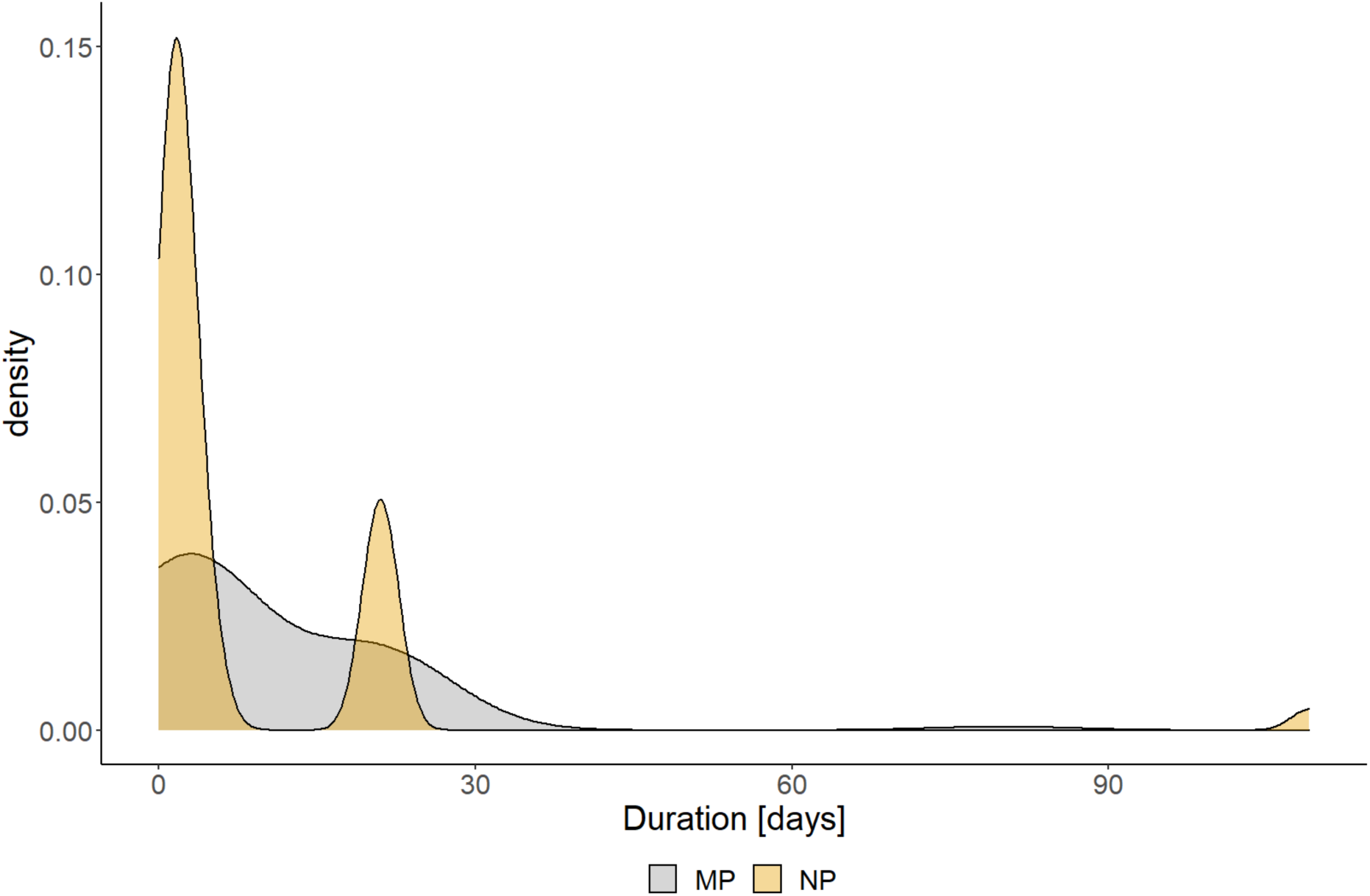
Density plot summarizing the duration of NMP experiments with *Daphnia* in days. NP (orange) experiments most often followed the recommendations of OECD guidelines 202 (48h)[27]and 211 (21 days)[28], while in MP (grey) experiments, modifications occurred.

##### 3.2.2.2 Adaptions and recommendations for standard laboratory procedures in NMP testing

To address the specific properties of NMP, several modifications to standard laboratory protocols have been proposed as standard requirements[107]. These include procedures to avoid NMP contamination via laboratory processes, the use of appropriate reference materials (e.g., particulate reference materials like naturally occurring particles) and adequate particle-free controls (e.g. including the same amount of solubilizers (Tween) as in NMP treatments). These measures are required to achieve construct validity by ensuring that we measure what we want to measure (i.e., the impact of the focus particles compared to other particles and a particle-free control) while avoiding contaminations in control treatments. Further, Nasser & Lynch (2019) recommend to pre-condition particles in medium containing natural organic matter (e.g., medium in which *Daphnia* have been kept for some hours), which leads to the formation of an ecocorona on the particles, which can affect toxicity outcomes (we discuss this further in section “3.3.4.2 Surface modifications ” below)[108].

In addition, an adaption of the OECD 202 guideline (acute test) for particle-based toxicant exposure has been proposed by Baumann et al. (2014), recommending an extension of the test period to 96h, since nanomaterials tend to show effects only after extended exposure[109].

### 3.2 NMP properties

#### 3.3.1 Polymer type

In total, 20 different polymer types were used in the screened literature (Fig. 4). Overall, the predominant polymer types were PS (51 %), PE (18 %), and PET (5 %). The diversity of polymers used in the MP studies was considerably higher than in NP studies (Fig. 4). Almost all NP studies used PS (90%), whereas the proportion of MP experiments conducted with PS was 33 %. The second and third most used polymer types in MP studies were PE (27 %) and PET (7 %).

**Figure 4.**
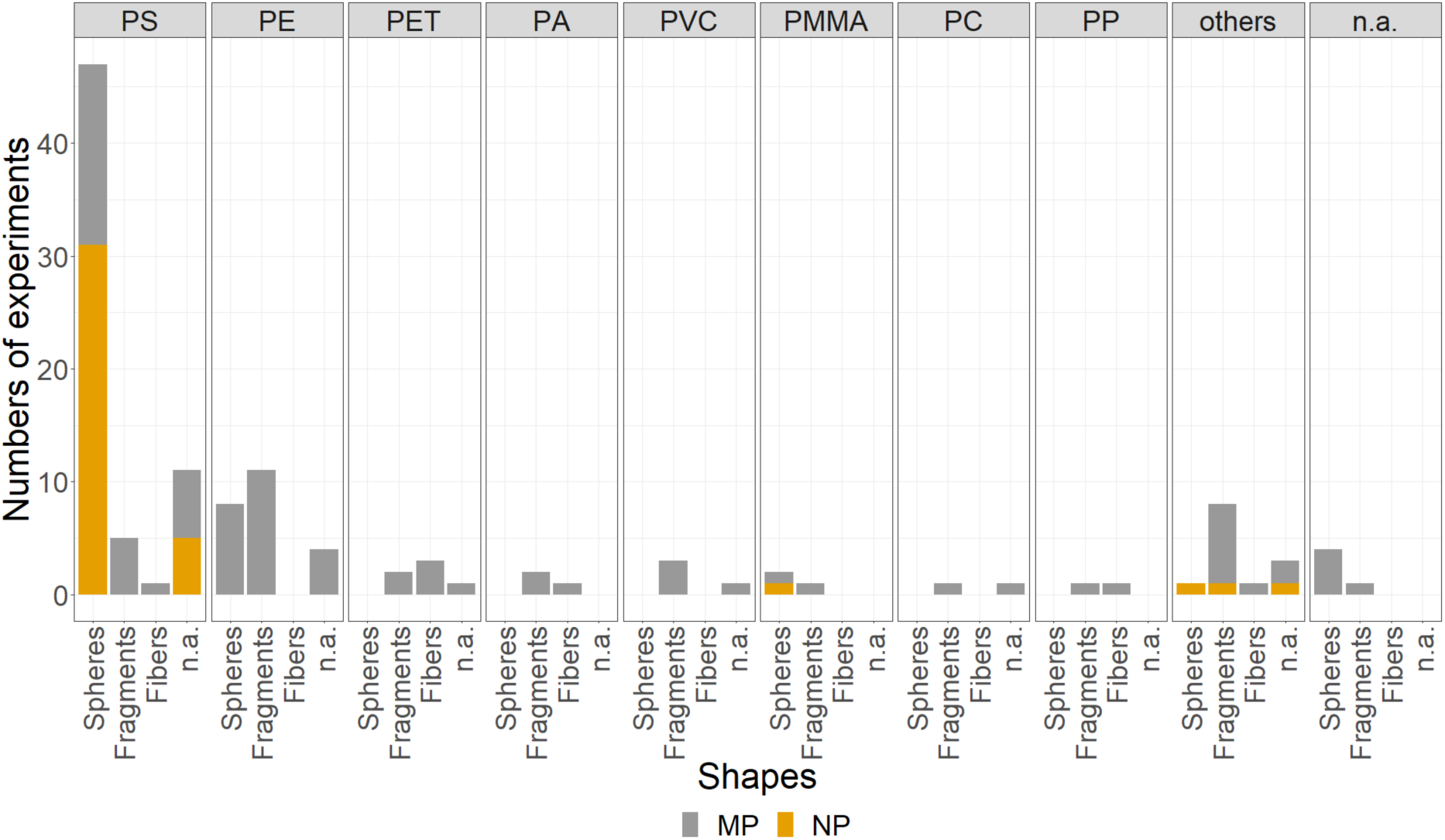
Number of different polymer types and shapes used in NMP experiments with *Daphnia*. Data is separated for MP (grey) and NP (orange). The category others contain Ethylene acrylic acid copolymer (EAAC), Polyhydroxybutyrat (PHB), Acrylnitril-Butadien-Styrol-Copolymer (ABS), Ethylene-vinyl acetate copolymer (EVA), Polybutylenterephthalat (PBT), “smoked cigarette buds” (probably cellulose acetate) and tire particles. Abbreviations: n.a.: polymer type or shape not reported.

A wide variety of polymers is used in industry, with PP, LDPE, PET, HDPE, PVC, PUR and PS being the most used ones in Europe[110]. In accordance, the polymers most commonly found in environmental samples of freshwater environments are also PE (24 %), PP (24%), PS (13 %), and PET (11 %)[7,111]. The almost exclusive use of PS in experiments with NP (90% of experiments) does not reflect this variety in polymer types found in the environment. The choice of polymers was more balanced in experiments with MP, but PET and PP were strongly underrepresented compared to their occurrence in environmental samples (5 MP experiments with PET; 2 MP experiments with PP). A likely explanation for the strong focus on PS is that NMP particles of this polymer are commercially available for scientists in specific size ranges and shapes. However, it has been shown that even supposedly identical 3µm PS beads from different manufacturers substantially differ in their material properties influencing particle-cell interactions and cellular responses[112]. This fact underlines that in-depth NMP characterization is needed to obtain comparable results in effect studies. Manufacturing non-commercially available NMP suitable for direct use in experiments with *Daphnia* is challenging and requires expertise in polymer fabrication and the development of novel methodologies.

Polymer types may differ in their effects on organisms [67,113]. Whether the observed differences between the tested NMP can be attributed solely to the polymer type remains however questionable, as none of the publications compared particles that differed only in polymer type while keeping all other NMP properties identical[112]. There is evidence however for other organisms that the polymer type may play a crucial role in determining toxicity (for example,[113]). Following similar approaches would thus be promising for understanding the polymer-dependent toxicity of NMP in *Daphnia*.

##### 3.3.1.1 Density

The density of NMPs is directly linked to their buoyancy and is primarily determined by the polymer type. When conducting experiments with pristine particles, density thus is the main driver of buoyancy irrespective of the NMP’s size[114]. Based on their buoyancy in water, polymers can be categorized into three groups: positively buoyant polymers (i.e. PE (density 0.89-0.98 g/cm^3^) and PP (density 0.85-0.92 g/cm^3^)), neutrally buoyant (i.e. PS (density 1.02-1.08 g/cm^3^)) and negatively buoyant polymers (i.e. PVC (density 1.30-1.58 g/cm^3^) and PET (density 1.29-1.40 g/cm^3^))[115]. In the studies screened, we found the use of 20% positively, 56 % neutrally, and 24% negatively buoyant MNPs (MP: positively (29%), neutral (39%), and negatively buoyant (32%); NP: positively (3%), neutral (92%), and negatively buoyant (5%)). As NP studies almost entirely used PS particles, most studies used neutrally buoyant particles, whereas the buoyancy distribution for MP was broader.

Pristine particles applied under controlled laboratory conditions without perturbation usually accumulate in the water layers according to their density[116]. Encountering rates of different *Daphnia* spp. with these particles may thus differ, if pristine particles are used together with comparably large or deep test vessels. Further, the jar volumes differ between different experimental setups, which may cause differing encountering rates. For example, in the OECD Test Guideline 202 a volume of 2 ml/animal is recommended[27], while in the OECD Test Guideline 211 ml a volume of 50 – 100 ml/animal is recommended[28].

In addition, while most *Daphnia* species are strictly limited to filter-feeding in the water column, *D. magna* often dwell near the ground and stir up particles and food material from the substrate. Compared to other *Daphnia* spp., individuals of *D. magna* might thus face particularly high concentrations and a higher risk of being affected by negatively buoyant particles, both in the lab and the natural environment.

#### 3.3.2. Size

The screened studies used particle sizes ranging from 1-366 µm (mean size) for MP studies and 20-800 nm for NP studies (Fig. 5).

**Figure 5.**
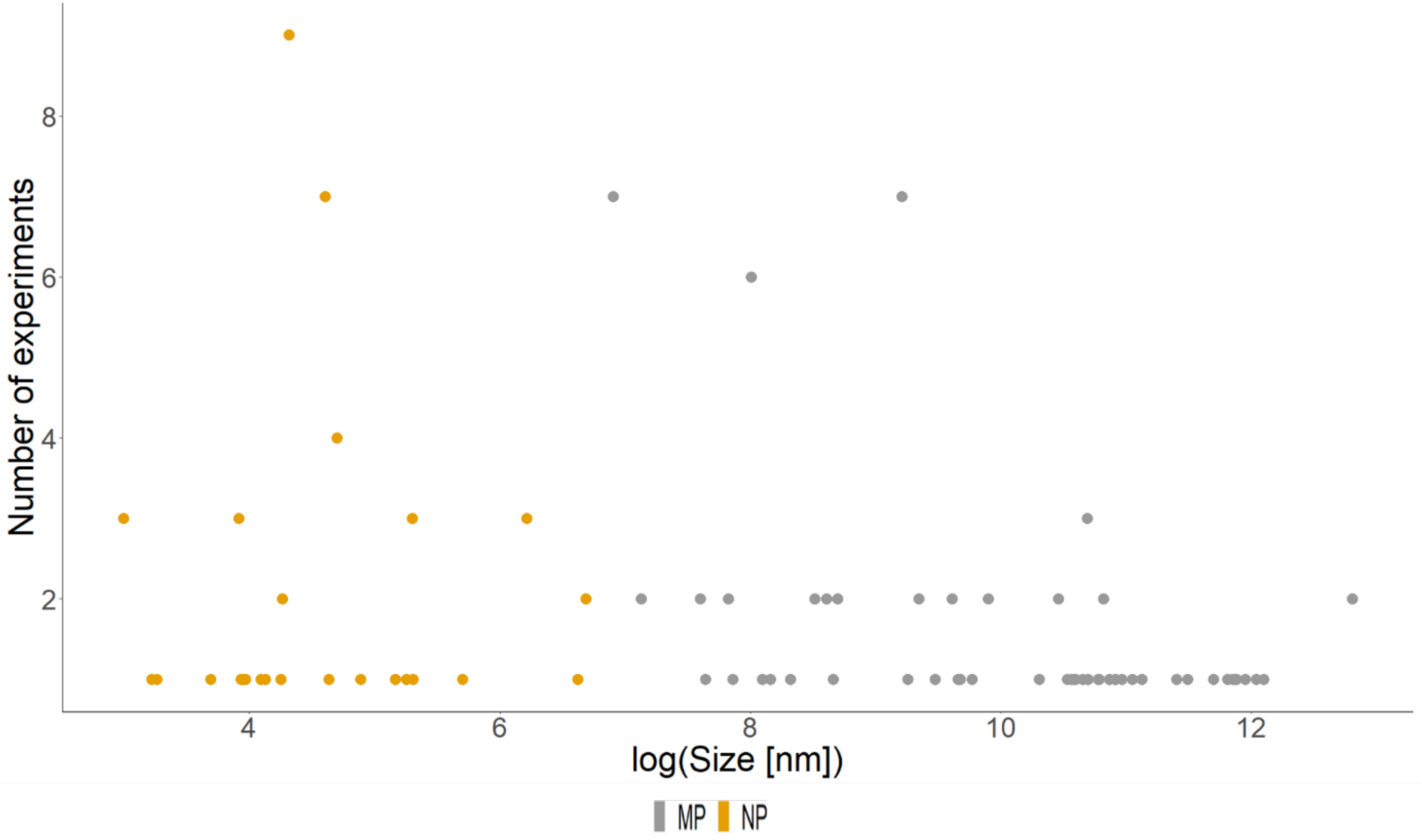
The density of the log(Size) in nm of used NMP particles in the screened experiments. The blue dashed line indicates the threshold between MP and NP particles at 1µm.

**Figure 6.**
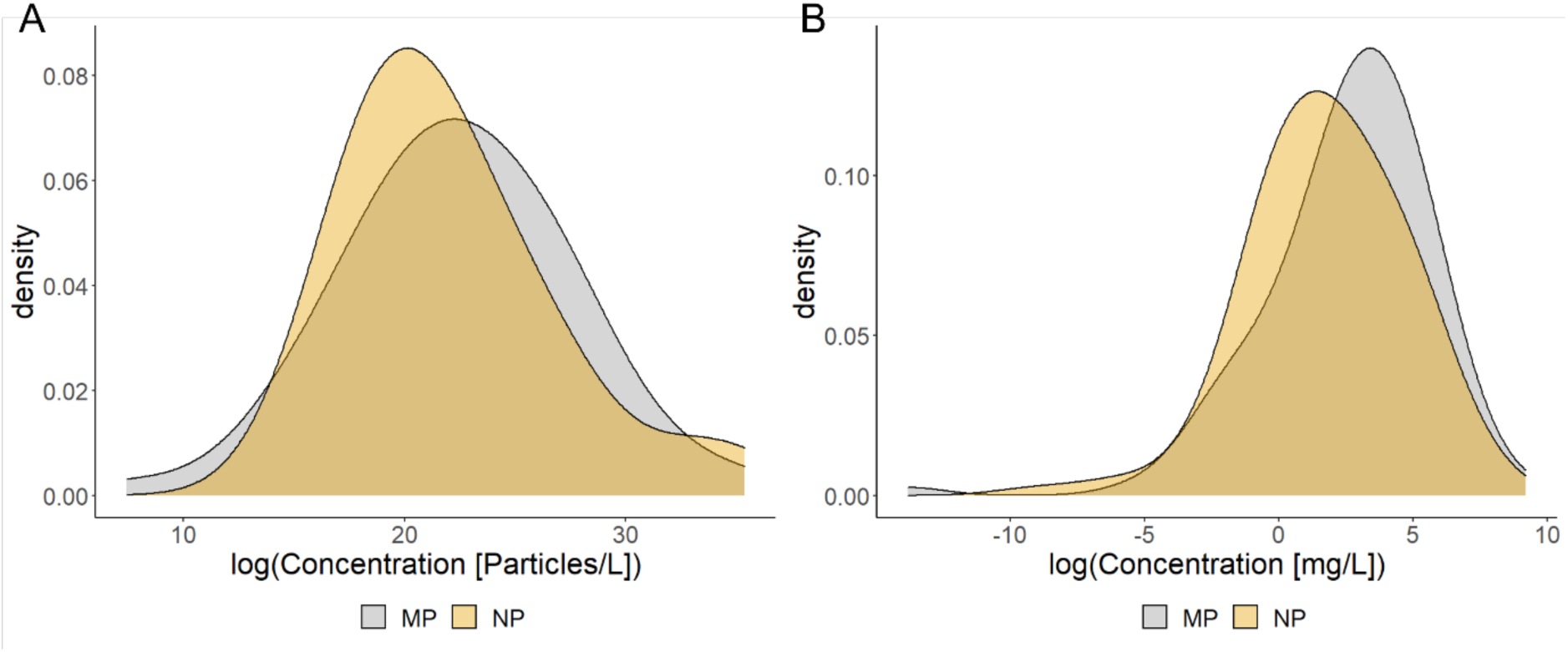
Density plots summarizing the concentrations in NMP experiments with *Daphnia*. (A) Density plot, showing log(concentration) of NMP given in particles/L. The data is separated for MP (grey) and NP (orange); (B) Density plot, showing log(concentration) of NMP given in mg/L. The data is separated for MP (grey) and NP (orange).

In contrast to information on the polymer type, information on the exact size range of NMP found in the environment are scarce. This is amongst others because universally accepted quantification and qualification tools of freshwater microplastics are lacking, and most studies use nets with mesh sizes larger than 300 µm to sample MP in freshwaters[117].

However, recent studies performed standardized sampling down to 10µm by the use of special filtering devices and subsequent-FTIR analysis[118,119]. As NMP size is a key property influencing NMP toxicity in *Daphnia*, standardized sampling methods for even smaller particles would be beneficial to reliably estimate NMP risk.

*Daphnia* usually ingest food particles between 700nm[120] and 70µm[121] depending on species and age. The filter apparatus usually does not catch particles smaller than this range, and larger particles are too large to enter the digestive tract. In general, smaller particles are considered more harmful than larger ones. For instance, An et al. (2021) found that the adverse effects of PE fragments (size classes: 17.23µm and 34.43µm) were more pronounced for smaller than for larger MPs [122]. Similarly, Schwarzer et al. (2022) found more substantial toxicity for smaller than for larger particles for both PS fragments (size classes: 5.7µm and 17.7µm) and spheres (size classes: 6µm and 20 µm)[123].

An aspect linked to the size-dependent effects is the potential of translocation through biological barriers, for example, from the digestive tract into tissues. Unfortunately, to date, it is still difficult to verify the translocation of NMP into tissues due to cross-contamination issues or movement of particles during sectioning of histological samples. Regarding translocation into the gut tissue or adhesion to the gut epithelium, it must be mentioned that *Daphnia*, like most arthropods, possess a so-called peritrophic membrane, which coats the midgut[124–126]. The peritrophic membrane may inhibit bacterial colonization of the gut epithelium, protects the gut epithelium from damage through particles, and limit particle translocation into the gut tissue to particles smaller than 130 nm in *D. magna*[127]. Hence, the pore size of the peritrophic membrane, which may differ between *Daphnia* species, has to be considered in all experiments testing the toxic effects of NMPs. This is an important aspect rarely considered in toxicity studies with *Daphnia* using particulate contaminants.

#### 3.3.3 Shape

The dominant shape of NMP used in the screened literature was spherical (49%), followed by irregular (28%) and fibrous (6%) (Fig. 4). However, 17% of studies did not specify the shape of the used particles. Research conducted with NP predominantly used spherical particles (82%) (16% no specification). For research with MP, the dominant shapes were irregular (40%) and spherical (35%), while only a few used fibers (8%) (17% with no specification).

This displays a certain misbalance to the shapes found in nature. For example, secondary MPs like fragments and fibers are much more common than spherical beads in freshwater environmental samples[8,111,128–131]. The focus on spherically shaped beads may result from their commercial availability. Some researchers tackle this issue by establishing protocols for the self-made production of fibers and fragments[123,132]. Although obtaining NMP in different shapes, size classes, and amounts necessary for large experimental setups is complex, it would be necessary to include more environmentally relevant particle shapes in toxicity testing.

The shape has been shown to directly affect the outcomes of *Daphnia* NMP toxicity tests. However, to test for the effect of different shapes, predominantly MPs of different polymer types were used[133,134]. These approaches cannot determine whether the observed effects are polymer or shape-dependent. To overcome this issue, Schwarzer et al. (2021) used three shapes (fibers, fragments, spheres) of the same polymer (PS) and showed shape specific responses in life-history traits and the morphology of *D. magna* after chronic exposure[123]. However, as for the polymer type, concluding the effects of a specific shape remains challenging, as often more than one NMP property is altered in comparative *Daphnia* effect studies.

Mechanisms leading to the observed differences are, up to now, not completely understood but likely include differences regarding residence times in the gut, clearance rates, damaging of gut tissue, clogging of the digestive tract or alterations of the gut microbiome [135–138]. For example, fragments with sharp edges are more likely to produce fissures in the peritrophic membrane, increasing the likelihood of NMP tissue translocation[139]. In addition, fibers might adhere to the gut epithelium and affect nutrient uptake adversely due to gut clogging[140]. Therefore, even if NMP particles of an, e.g., fiber shape, might not inflict direct physical damage on the epithelium of the midgut, effects on another level may be detectable[141].

#### 3.3.4 Additives

For most NMP particles, no information about the additives incorporated in the used polymer were provided (74% overall; MP 62%; NP 95%). When provided, the most frequently described additives were colour pigments (9%), followed by benzophenone-3 (BP-3) (3 MP experiments). However, most often, the information on the additives remained vague (for example describing that NMP contained minimum amounts of additives, contained (undefined) stabilizers or that additives were detected, but without further specification).

Almost all synthetic polymers include additives to give them their unique characteristics[142], like enhanced thermal stability or specific flexibility [143]. Plastic additives have been shown repeatedly to affect outcomes in toxicity tests, when leaching out of the polymer during exposure [144]. For example, Schrank et al. (2019) showed that the plasticizer DiNP leaches from flexible PVC into the surrounding media[145], causing toxic effects in *D. magna*[145,146]. As the composition and amount of additives often remain unreported and information on additives is not always available for commercially available NMP, results from NMP toxicity tests are often hard to interpret. The need thus arises to distinguish between the effects caused by the NMPs themselves and possible effects from associated additives[145]. One option is to characterize all compounds of the used NMP as shown for freshwater mussels[113]. Another approach is to compare the effects of particles with additives with the effects of particles without additives, but with otherwise similar properties (i.e., same size, shape, polymer type, etc.)[147]. Due to difficulties in manufacturing NMPs without additives, the experimental implementation, however, remains challenging.

##### 3.3.4.1 Fluorescence tags

Fluorescence tags are frequently used in NMP research to monitor the uptake of NMP in *Daphnia*[148–150]. Fluorescence tags were used in 33% (32% MP; 33% NP) of the screened studies, and 67% of these studies gave detailed information about the properties of these tags (name, color, excitation, and emission wavelengths).

The use of fluorescent particles might be problematic in studies investigating the effects of NMP and the translocation of particles into tissue. Similar to other additives, it has been shown that fluorescence tags can leach from the polymer particles, especially in alkaline aqueous media (which most standard media are), and that they may cause toxic effects[151–153]. Furthermore, fluorescent dye leaching from the particles into the surrounding tissue may be erroneously interpreted as particle tissue translocation[151]. Therefore, fluorescent labelled NMPs can be used for specific research questions but it should be considered that the environmental relevance of this particles is not given.

###### 3.3.4.2 Residual monomer contents

None of the publications we screened reported on the residual monomer content of NMPs. This is probably due to missing information from the manufacturers and the very high effort for the scientists to determine it. Nevertheless, residual monomers of the polymers might also have the potential to leach out, causing harm to organisms[154,155]. For example, the monomer of styrene has been described to be highly toxic to aquatic organisms, including *D. magna*[156]. Interestingly also supposedly identical MP particles (in this case, plain PS beads Ø 3µm from two manufacturers) can differ in their monomer contents[112], possibly reducing comparability between experiments. For scientist working with NMP in toxicological experiments, information on residual monomer contents can only be obtained through in-depth characterisation of the particles used. Methods that can be and have been applied are i.e. NMR-spectroscopy and GC-MS[112,113].

##### 3.3.4.3 Surface modifications

In the publications we screened, information on surface modifications was given for 59% of all used particles (53% MP; 78% NP). For 41% of particles, no information was given or the particles were unmodified.

###### 3.3.4.3.1 Uncontrolled surface modifications

Information on uncontrolled surface modification was given on 29% of particles used (28% MP; 31% NP). This included milling/grounding of particles (13% overall; 21% MP; 2% NP), sonification (none MP; 2% NP), UV-weathering (1% MP; none NP), exposure to the abiotic environment like the incubation in test medium, and the sorption/addition of organic substances like palmitic or humic acids (2% MP; 28% NP), and the exposure to biotic environment/wastewater (2% MP; none NP).

Milling[157], sonification[158] and artificial weathering under UV light[159] can substantially alter the properties of NMP. In the reviewed literature, we found that the distinction between milling and weathering was not always apparent. For example, some studies reported using “artificially weathered” particles, where particles that had only been milled. Although both processes, milling and UV-weathering, lead to the fragmentation of particles (i.e., after the process mean particle size is smaller), the change in NMP properties resulting from them is quite distinct: While milling, like sonification, mostly leads to changes in the particles’ surface structure and shape, but not to chemical changes[158,160], UV-weathering causes additional chemical changes (including changes in surface charge and composition of functional groups) and embrittlement due to photo-oxidation[15,159,161,162]. While the size of NMP particles and the molecular weight of polymer chains decrease with increasing exposure to UV radiation, the amount of carboxylic acids, peroxides and ketones on the surface increases, a pattern that seems to be generalizable across different polymer types (LDPE[159]; PP[163]; PS[15]).

None of the studies we screened tested explicitly for differences between milled, sonicated, or UV-weathered particles. One study compared pristine and UV-weathered MP, revealing no effects for both particle types[164], and one study compared sonicated to non-sonicated NP, revealing a twofold lower toxicity of the sonicated NP particles[165]. While there are hardly any effect studies with *Daphnia* and NMP focusing on altered surface properties, studies with murine[112] and algal[166] cells show that these can have a major influence on the toxicity of NMP.

The incubation of NMP in biotic environments like wastewater or natural lake water and the incubation in medium containing artificially added organic substances or any dissolved organic carbon can also lead to substantial changes in the particles’ surface structure. Organic substances attach to the particles’ surface, forming a so-called ecocorona which may change the particles’ stability, reactivity (interaction with the surrounding media or organisms) and toxicity[167]. Schür et al. (2021), for example, found that the pre-incubation in wastewater reduced the toxicity of milled irregular PS MP (<63µm) in comparison to the same but pristine MP on *D. magna*[168]. However, the picture may change for smaller sizes of NMP where tissue translocation may occur. For instance, in experiments with murine macrophage cells, it has been shown that PS beads (Ø 3µm) coated with an ecocorona attach more readily to the cell surface and are internalized more frequently than PS particles without ecocorona[39].

When particles are large enough bacteria start settling on the attached biomolecules and forming a biofilm[38]. The diversity of bacteria and the complexity of the biofilm depends on the particles’ initial surface charge, the surface smoothness and the origin of the used incubation medium (e.g., freshwater or saltwater)[169,170]. Biofilms lead to several changes in NMP properties, including changes in the particles’ surface properties, increased leaching of additives and residual monomers, increased embrittlement and loss of mechanical stability due to attacks of enzymes and radicals, the accumulation of water in the polymer matrix (causing swelling and increased conductivity) and the excretion of lipophilic microbial pigments[171]. As a result, biofilms can impact the effects of NMP on organisms in the environment. For example, in a recent study, Motiei et al. (2021) found that the toxicity of MP on *D. magna* was ameliorated in the presence of biofilms, and the authors hypothesized that the daphnids were able to digest biofilms as kind of an additional microbial food supply[172].

###### 3.3.4.3.2 Controlled surface modifications

Controlled surface modifications were reported for 30% (24% MP; 37% NP) of particles. These were: carboxylation (11% of all studies reviewed; 7% MP; 14% NP), amination (6% overall; 1% MP; 12 % NP), the coating with tensides like Tween, Triton X-100 or SDS (10% of all studies reviewed; 14% MP; 5% NP), and staining with Nile red, a dye that attaches to the NMP surface (2% overall; 1% MP; 2% NP).

As with uncontrolled surface modifications, there is little information on how controlled surface modifications influence NMP effects on *Daphnia.* However, due to changes in the sorption behavior of charged particles[173], differing effects on *Daphnia* appear likely. For instance, Zhang et al. (2020) compared the effects of iron-doped and non-iron-doped MP (PS spheres, Ø 1µm) with two different surface modifications (carboxylation, amination) and found that the aminified particles were, in general, more harmful than the carboxylated ones[174]. A similar trend has been found for gold nanoparticles [175], which underscores the importance of surface charge in *Daphnia* effect studies, at least for small particles.

#### 3.3.5 Concentration

All screened studies reported the NMP concentrations used in the experiments, but the units for describing the concentrations differed: while 28% of concentrations (32% MP; 9% NP) were given as particles per volume, 72% of concentrations (68% MP; 91% NP) were given as particle weight per volume (including ppm and mmol). Overall, concentrations were quite similar in MP and NP studies (Fig. 7).

**Figure 7.**
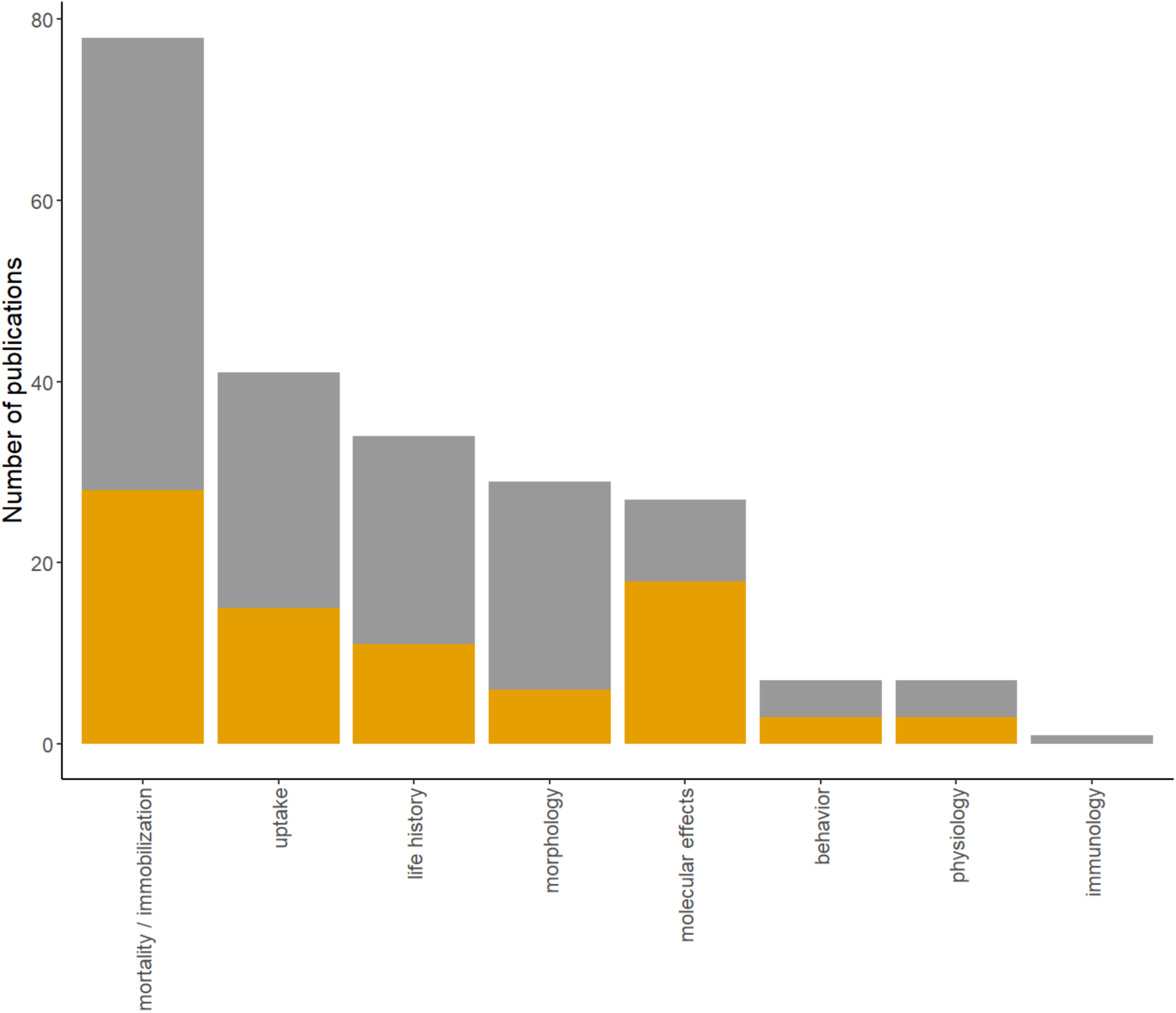
Barplot showing the analyzed endpoints in publications with NMP and *Daphnia*. Data is separated for MP (grey) and NP (orange); Endpoints were counted when mentioned in the text.

Most screened studies used MP concentrations much higher than those reported in the environment (for MP between 0.9 to 25.8 particles/L[176–178]). However, reliable information about environmentally relevant concentrations of smaller-sized NMP in the range ingested by *Daphnia* spp. is lacking due to analytical detection limits, and actual concentrations of these smaller particles may likely be higher than those reported for larger MPs[179]. Therefore, the estimation of the environmental risk that NMP impose on *Daphnia* spp. remains challenging. Considering that effects become more apparent at higher concentrations [180–182] and most NMP studies with *Daphnia* used high concentrations, the effects of NMP in today’s environment may be overestimated. Therefore, a focus on experiments with environmentally relevant concentrations would be necessary to understand the current effects of NMP on organisms[183]. In addition, focusing on dose-response experiments conducted with a range of NMP concentrations may provide more detailed knowledge about concentrations at which NMP might become problematic, if the amount of NMP increases in the environment, as expected.

## 4. Endpoints

As the specific endpoints analyzed for *Daphnia* in toxicity experiments are manifold, we focus on giving a brief overview of what has been analyzed and which parameters have been shown to affect the different endpoints. A detailed assessment of the different endpoints analyzed for *Daphnia* concerning MP toxicity can be found in the review by Yin et al. (2023)[184].

41% (42% MP, 38% NP) of reviewed articles analyzed NMP uptake (i.e., whether particles were found in the intestinal tract and/or number of ingested particles at a specific time point) or the uptake rate (i.e. the number of particles ingested over time; Fig. 7). Only a few studies (7 %) analyzed behavioral effects (6% MP; 8% NP) including both, feeding and swimming behavior. The physiological response of daphnids to NMP exposure was measured in 7% of reviewed publications (6% MP; 8% NP) which measured changes in the daphnids’ heartbeat rate. Effects on the molecular level were investigated in 27% of publications (15% MP; 46% NP) and included the analysis of gene expression (heat shock proteins, transcriptomics), proteomics, DNA damage (comet assays) and the amount of reactive oxygen species (ROS) including the ROS detoxifying enzymes (catalase, superoxide dismutase, glutathione-S-transferase) and the relevant molecular markers for ROS (e.g. malondialdehyde (MDA)). One study investigated the immunological response of *Daphnia* to NMP utilizing changes in haemocyte numbers. Morphological effects were investigated in 29% (37% MP; 15% NP) of publications, including measurements of body length, relative body width and tail spine length. Effects on the life history (including reproductive output) were investigated in 34% (37% MP; 28% NP) of publications and included the number of offspring, time to first brood, and survival/fitness of the offspring. Mortality and immobilization have been investigated in 77% of studies (81% MP; 72% NP) and thus most frequently in the reviewed literature.

Understanding the mechanisms that lead to toxic effects in *Daphnia* individuals and populations requires investigating a broad range of endpoints. By combining the results along effect cascades, adverse outcome pathways (AOPs; [185,186]) can be established, linking NMP uptake to molecular and individual level effects (behavioral changes, physiological response, immunological response), to effects on reproduction and offspring (offspring numbers, morphological alterations, production of males), and finally the population level (e.g. *Daphnia* abundance). In addition, it might be helpful to identify particularly sensitive endpoints to compare the toxicity among NMPs with different property combinations and identify those traits or trait combinations associated with toxicity.

### Uptake

Uptake links the exposure concentration in the medium to the actual dose a daphnid has to deal with in experiments. It thus marks the starting point of an AOP and the transition from exposure to hazard assessment. Fluorescently labeled particles are usually used in experiments to monitor and quantify the uptake of particles. However, as discussed above, when effects are to be analyzed in parallel, control treatments to test for the potential toxicity of the fluorescent labels should be included.

### 4.1 Behavior

When exposed to NMP in the medium, several comparably quick responses are possible: First, mere contact with non-digestible particles may change the daphnids’ feeding or swimming behavior. For NMP exposure however, no consistent trends in changes of the swimming behavior can be identified. On the one hand, exposure to NMPs led to a reduction in swimming activity in *D. magna* (PE fragments (48 µm) [187]; PS spheres (110 nm)[188] & PS NP agglomerated/non-agglomerated[165]). On the other hand, there are studies, showing an increase (PS, probably spherical since particles were purchased (1 & 10 µm)[189]) or no impact (PS-NP (20 & 200 nm)[190]) on the swimming behavior of *Daphnia*. The conflicting results may indicate that swimming activity is not an optimal endpoint to elucidate the effects of NMP on *Daphnia*.

In regards to feeding behavior, for example Chen et al. (2020)[191] reported an increase in filtration activity at high MP concentrations (PS spheres (5 µm)). A similar trend was observed by Hoffschröer et al. (2021), where the ingestion rates of MP (PS spheres (1 µm)) were increased in conditions of low food and high temperatures[102]. They further concluded, that the observed trends, reflected the complex regulation patterns of the water current generated by the animals’ thoracic limbs[102].

As pointed out by Bownik (2017)[192], the analysis of the behavior of *Daphnia* covers an extensive range of possibilities (e.g. feeding/filtration, swimming distance, velocity, sinking rate, swarming) which can be useful for hazard assessment, especially when connected to automated data acquisition to reduce observer bias. In studies dealing with NMP, only a few of these parameters have been assessed, opening new possibilities for future studies.

### 4.2 Physiology

Second, NMP exposure and uptake may lead to changes in the daphnids’ physiology, like metabolic or respiration rates. The only physiological response measured in the reviewed literature was the daphnids’ heartbeat rate. Here, Xu et al. (2020) showed that exposure to primary PS-NP (20 nm) reduced *D. magna* heartbeat rate [193]. Heartbeat rates typically assessed in drug testing and automatized procedures including blood flow measurements have been proposed [194–196]. Similar approaches may be equally promising for NMP studies, although differences seem to exist among different *Daphnia* species, with *D. magna* heartbeat rates being more robust than rates in other species[196].

### 4.3 Molecular and immunological responses

Third, the uptake of NMP can lead to molecular and immunological responses. Molecular effects include, among others, changes in gene expression (transcriptome), protein composition (proteome) or, more specifically, an increase in ROS enzymes. Immunological effects can help bridge the gap between molecular effects and those on the organismic level and thus contribute substantially to our understanding of mechanisms leading to toxic effects. An additional option to the targeted analysis of immune system specific gene expression[197] is to count the number of hemocytes in the hemolymph. Hemocytes are circulating phagocytic immune cells that perform the immune response in invertebrates, comparable to macrophages in vertebrates[198]. In the only study of the screened literature investigating the immune response, Sadler et al. (2019) showed that the number of haemocytes in *D. magna* was generally higher after exposure to NMP, and for some clones, this effect was more pronounced at higher temperatures[66].

### 4.4 Morphology

Changes on a molecular level can finally lead to effects on the organismic level. During development and reproduction, effects on *Daphnia* morphology can become apparent. For example, Schwarzer et al. (2022)[123] found a reduced body length of *D. magna* when exposed to small PS fragments and beads (5.7 µm and 6 µm, respectively). On the other hand, Liu et al. (2020)[149] found no impact on body length in *D. pulex* when exposed to PS-NP (75 nm) but on growth rate over generations. This indicates that even if morphological effects are not seen within individuals, effects on this endpoint might be detectable on a population level. *Daphnia* spp. can exhibit phenotypic plasticity in response to altered environmental conditions and adjust their body parameters (e.g., body length, body width, tail spine length) and life-history to changes in food concentration[199], food quality[87,200], and crowding factors[199,201]. The careful control of these parameters in toxicity experiments, including NMP studies, is thus required to avoid confounding effects.

### 4.5 Life history

The analysis of changes in life history is another well-established endpoint for organismic effects in *Daphnia* spp. and directly impacts reproductive output if, for example, maturation is delayed[28]. Life history traits that are often assessed in *Daphnia* spp. include the time until the production of the first clutch, time to primiparity (i.e., the release of the first clutch from the brood pouch), times between clutches, and time to death. The reproductive output in *Daphnia* is usually measured by counting offspring numbers per clutch or the total number of offspring at a specific time of exposure (usually after 21 days[28]). Bosker et al. (2019) [202] for example counted the offspring over generations to estimate the effects on *Daphnia*’s population and found a significant decline in *D. magna* biomass when exposed to PS beads (1-5 µm). Similar results were found by Aljabiachi et al. (2020)[104], who analyzed the effects of PS MP (15 µm) in a mesocosm experiment. For studies analyzing NP, effects like, e.g. reduction in the number of offspring and age at first brood have been found for PS-NP (90 nm)[148]. On the other hand, Rist et al. (2017) found no adverse effects on the number of offspring when exposing *D. magna* to MP and NP (2 µm and 100 nm, respectively)[203]. Besseling et al. (2014) used the number of offspring of *D. magna* and the size of neonates to analyze the effect of PS-NP (70nm, beads) and found a negative correlation between both life-history parameters and the concentration of NP[204].

These often contrasting results emphasize that effects on *Daphnia* are dependent on a mixture of specific characteristics of the performed experiment. Whether an effect of NMP on a specific endpoint elicits or not, may depend on three different levels, as depicted in the mindmap (Fig. 1). This variety of parameters may explain the variance in the reported effects. To be able to classify/interpret the effects of NMP, the pure consideration of an endpoint (effect yes/no) is therefore not meaningful, if one ignores the circumstances that may lead to effects.

### 4.5 Immobilization

Immobilization (and mortality) was the endpoint most frequently tested in the reviewed literature. This is reasonable, as immobilization is the standard endpoint used in acute toxicity testing[27]. The effects of NMP on *Daphnia* spp. mobility and survival and how these effects are modulated by parameters of *Daphnia* ecology, experimental parameters and NMP characteristics will be discussed through the meta-analysis below. As not all studies discriminated between immobilization (the organism does not move within 15 seconds after agitation) and mortality (absence of heart beat), we will refer to this endpoint as immobilization only.

## 5. Meta-analysis

For the meta-analysis of immobilization caused by NMP exposure, we extracted data from 59 studies resulting in 720 immobilization risk measurements. For 26 samples, concentrations were reported as parts per million. For another four samples, transforming particles per ml to mg per ml was impossible. The following meta-analysis is thus based on 690 data points.

Overall, the risk of immobilization was higher in treatments with NMP than in particle-free controls (most data points above the horizontal line in Figures 8-16).

**Figure 8.**
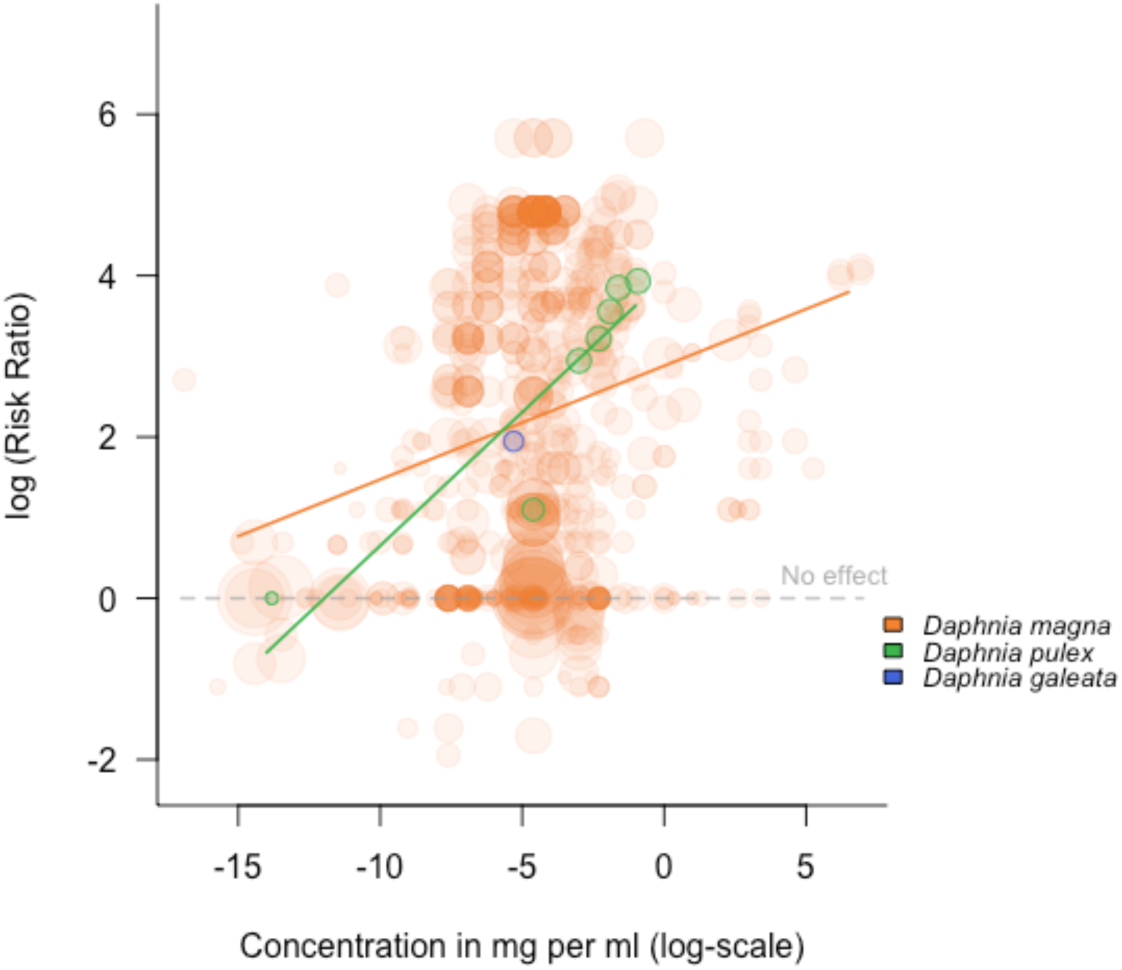
Log(RR) of immobilization rates of three different *Daphnia* species exposed to NMPs at different concentrations. Point sizes reflect the inverse standard error of the data points. Meta-regression lines are shown for groups with at least five data points.

### 5.1 Daphnia biology/organismic level

#### 5.1.1 Species

We found immobilization data for three *Daphnia* species: *D. magna* (54 studies contributing 682 data points), *D. pulex* (2 studies contributing 7 data points) and *D. galeata* (1 study contributing 1 data point). The mean risk of immobilization in response to NMP did not differ substantially between these three species but was slightly higher for *D. pulex* than for *D. magna* (see green regression line in Fig. 8 positioned above the orange line).

#### 5.1.2. Age and sex

Most daphnids used in immobilization tests were younger than 24 hours (595 data points). The oldest tested adults were 18 days old at the start of exposure. We found no apparent correlation between the risk of immobilization and the age of the test organisms at the start of exposure (Fig. S4). All studies included in the meta-analysis used females in their experiments.

### 5.2 Experimental setup

#### 5.2.1 Food

29 studies (contributing 209 data points) included in our meta-analysis provided food simultaneously to the exposure to NMP. Conversely, 19 studies (398 data points) did not provide food. In general, immobilization RRs were higher for experiments not providing food (see green regression line in Fig. 9 positioned above the blue line). This is particularly interesting, as food is usually provided in chronic experimental setups where we would expect effects to be stronger. Providing food during exposure thus seems to reduce the adverse effects imposed on daphnids by NMP.

**Figure 9.**
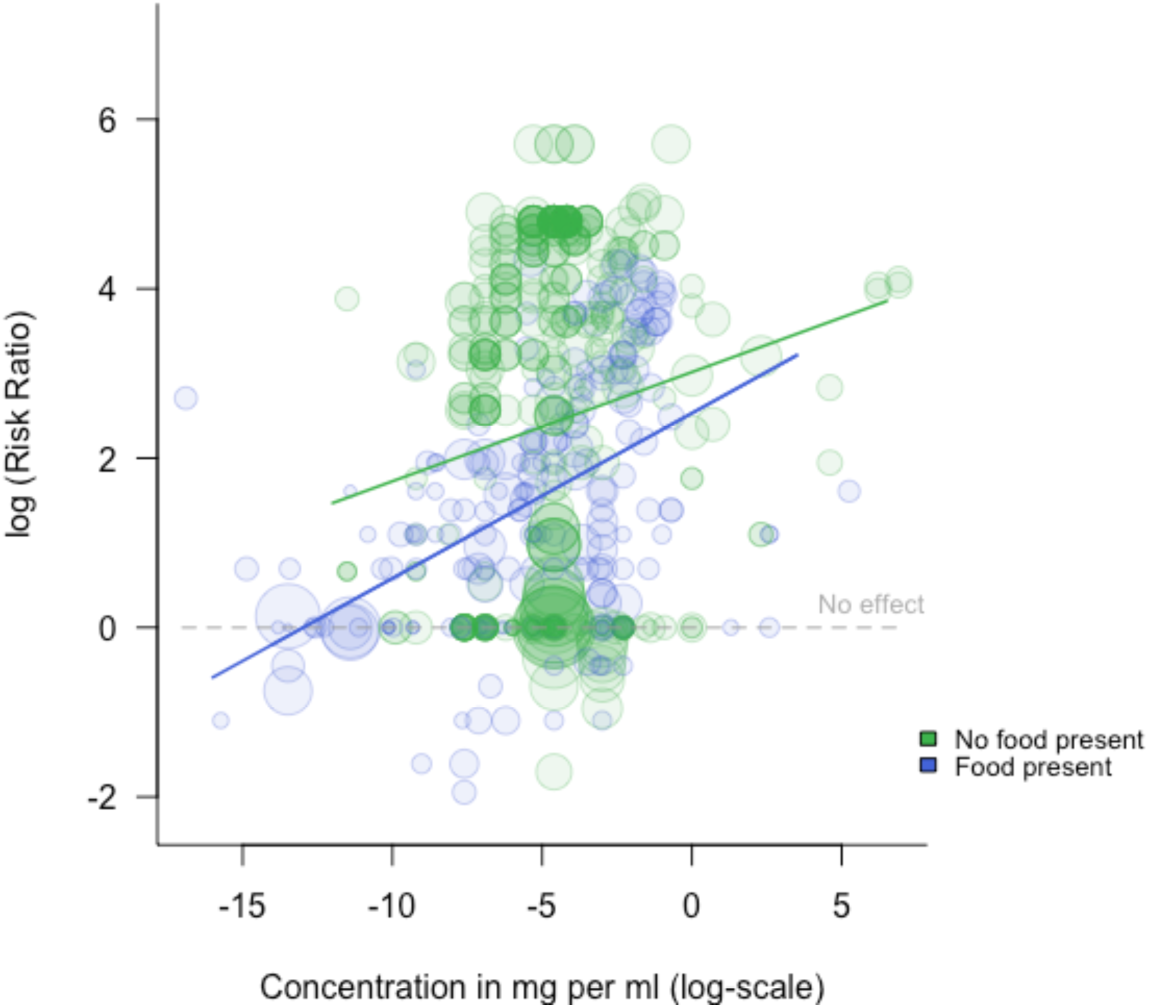
Log(RR) of immobilization rates of *Daphnia* spp. exposed to NMPs at different concentrations with (Food present (blue)) and without (No food present (green)) simultaneous provisioning of food. Point sizes reflect the inverse standard error of the data points.

#### 5.2.2 Temperature

The experimental temperatures in studies investigating immobilization ranged from 15°C to 30 °C. Immobilization RRs were generally slightly increasing with increasing experimental temperatures (Fig. S5).

#### 5.2.3 Exposure duration

All studies included in the meta-analysis were laboratory studies. They covered a wide range of exposure durations from 24 hours to 63 days (Fig. 10). Based on the meta-analysis dataset, immobilization risk seems to decrease with increasing exposure duration. For example, the immobilization risk was higher for individuals exposed for 48 hours than for those exposed for 21 days (see the light green regression line (48h) positioned above the orange line (21 days) in Fig. 10). However, short-term experiments were usually conducted with higher NMP concentrations. In contrast, chronic exposure experiments usually provided food and were more often conducted with lower NMP concentrations (see Fig. 10: NMP concentrations were, on average higher for 48-h than for 21 days of exposure). The decline in immobilization risk with increasing exposure time may thus likely be a result of confounding.

**Figure 10.**
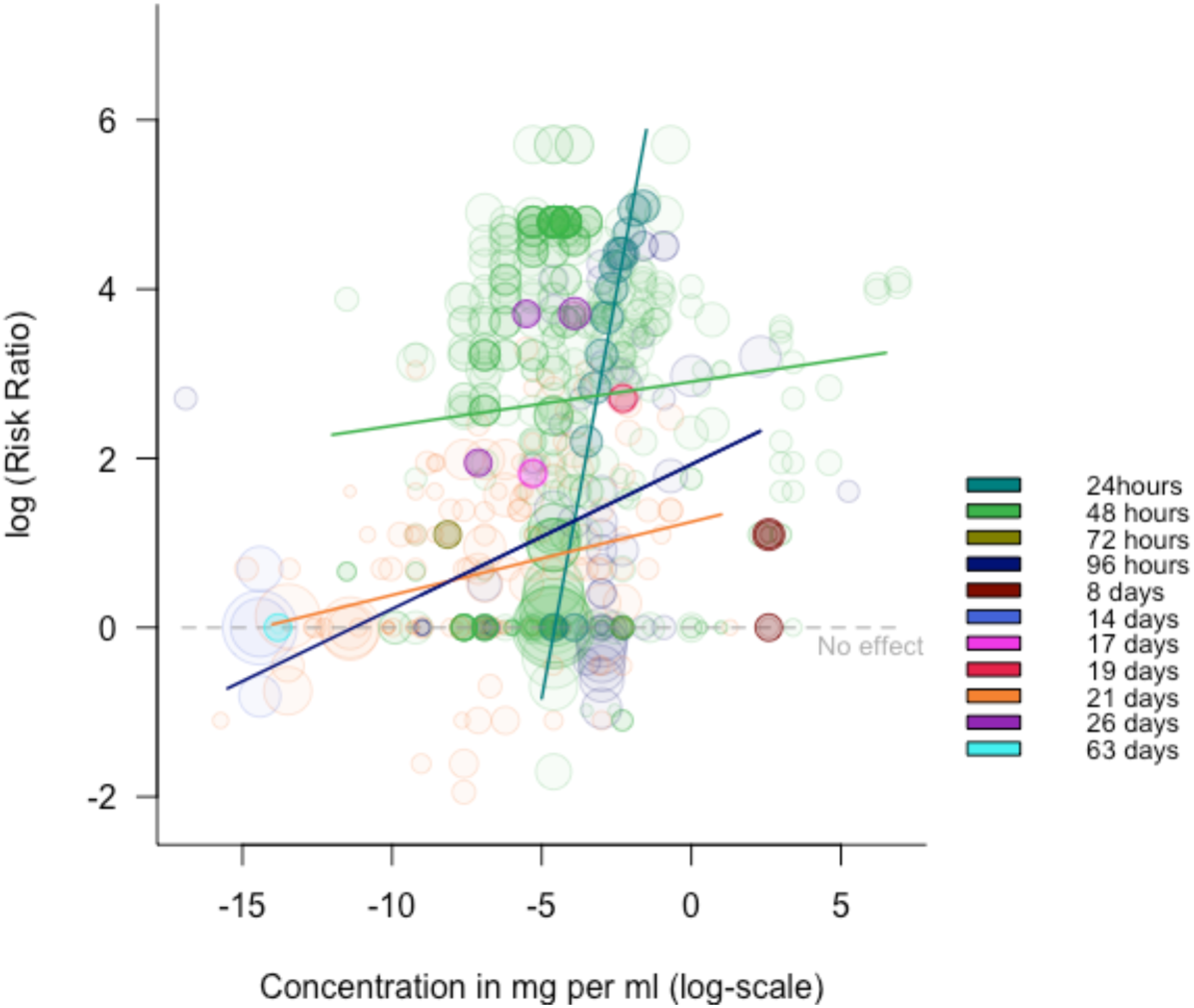
Log(Risk Ratios) of immobilization rates of *Daphnia* spp. exposed to NMPs at different concentrations for different periods. Point sizes reflect the inverse standard error of the data points. Meta-regression lines are shown for groups with at least five data points.

### 5.3 NMP properties

#### 5.3.1 Polymer type

In our meta-analysis, 17 different polymer types, including mixtures, were included (Fig. 11; see Table S1 for a complete list, including sample sizes). Different polymer types seem to be associated with different immobilization risks. For example, polylactic acid (PLA) was consistently more toxic than polyurethane (PUR). In addition, the concentration-dependent increase in effect size also differed among different polymers (see different slopes in Fig. 11): For example, while immobilization risk ratios in experiments with PS, PMMA and PMMA-PSMA increased only slightly with increasing concentrations, the concentration-dependent increase of toxicity was much steeper in PP, PLA and PUR.

**Figure 11.**
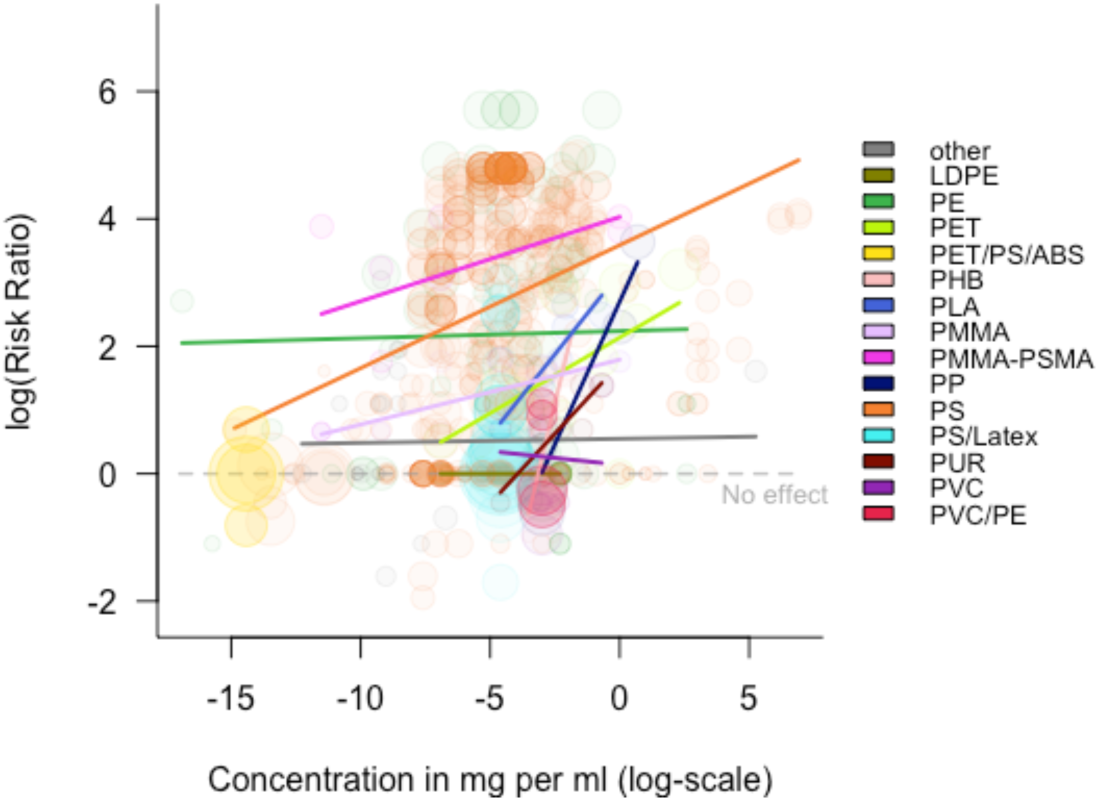
Log(Risk Ratios) of immobilization rates of *Daphnia* spp. exposed to different polymer types at different NMP concentrations. Point sizes reflect log inverse standard errors of the data points. Meta-regression lines are shown for groups with at least five data points. For a list of polymer names and sample sizes, see Table S1 in the supplementary online material.

#### 5.3.2 Size

In our meta-analysis dataset, 38 studies used MP contributing 274 data points, and 23 studies used NP contributing 416 data points. Immobilization risk ratios generally decreased with increasing particle sizes (Fig. 12).

**Figure 12.**
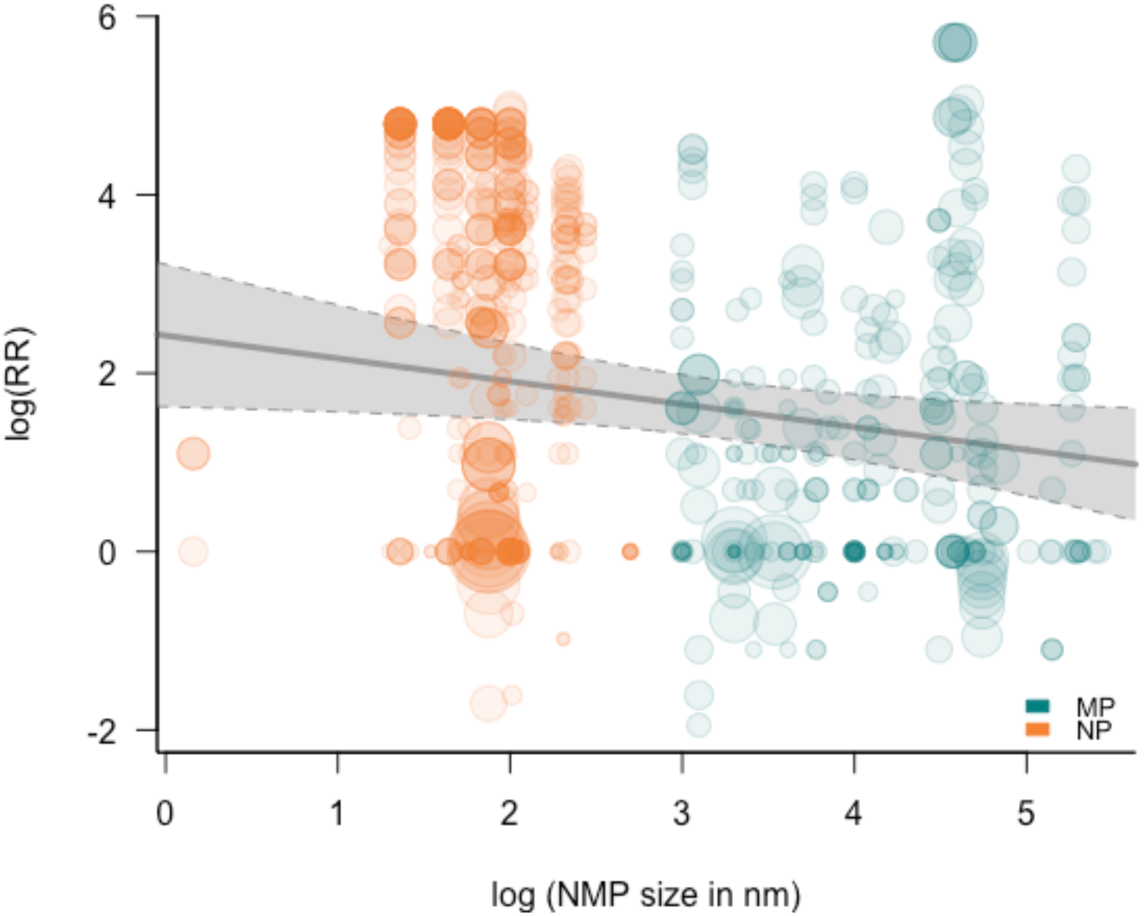
Influence of particle size on log(Risk Ratio) of immobilization in *Daphnia* spp. exposed to NMP. Point sizes reflect log inverse standard errors of the data points. The line represents the meta-regression line, and the shaded area represents the 95% confidence interval.

#### 5.3.3 Shape

In our meta-analysis, 42 studies investigated spherical NMPs (546 data points), 17 investigated irregular fragments (115 data points), 5 studies used fibres (11 data points), and one study investigated NP aggregates (6 data points). Fragments and spherical NMPs did not differ in immobilization risk ratios (Fig. 13), and the data for fibres and NP aggregates was too limited to draw any conclusions.

**Figure 13.**
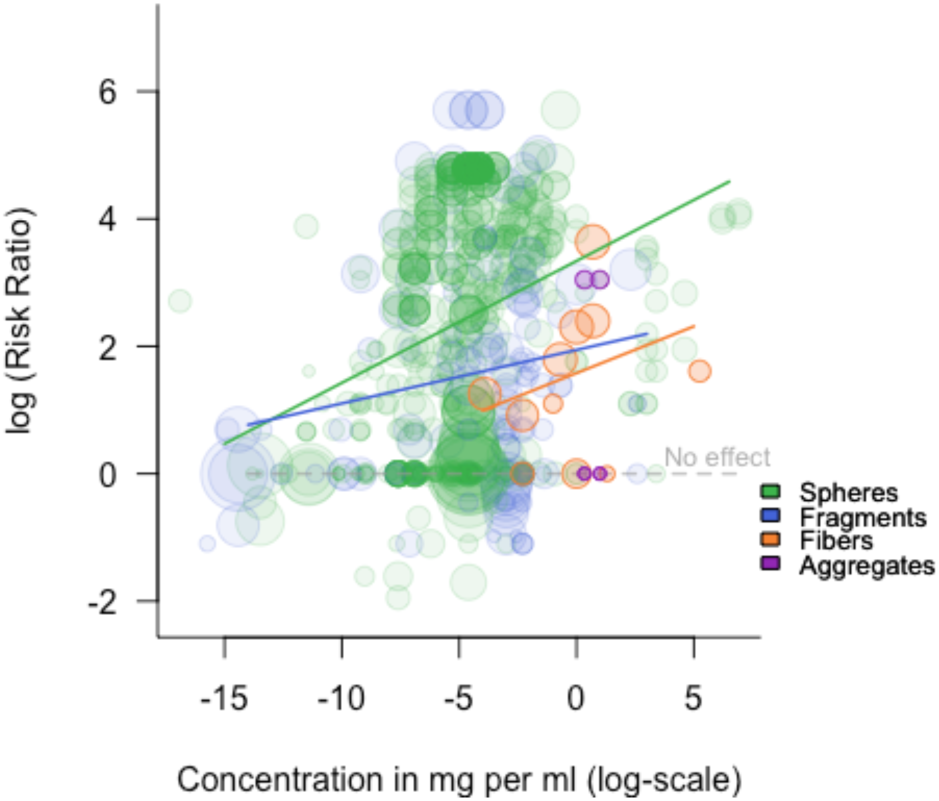
Log(Risk Ratios) of immobilization rates of daphnids exposed to NMP with different shapes at different concentrations. Point sizes reflect the inverse standard error of the data points. Meta-regression lines are shown for groups with at least five data points.

#### 5.3.4 Additives, fluorescence tags and other polymer modifications

Two studies in our meta-analysis used NMPs modified with the additive benzophenone-3 (BP-3, melt-blended; 8 data points), and one used 13C-labelled polyethylene (3 data points). 18 studies (contributing 131 data points) used NMP with fluorescence tags.

NMP containing benzophenone-3 increased immobilization rates compared to NMP with other chemical modifications or non-modified NMPs (see pink line in Fig. 14 positioned above other lines). According to the compiled dataset, the presence of fluorescence tags slightly decreased the risk of immobilization.

**Figure 14.**
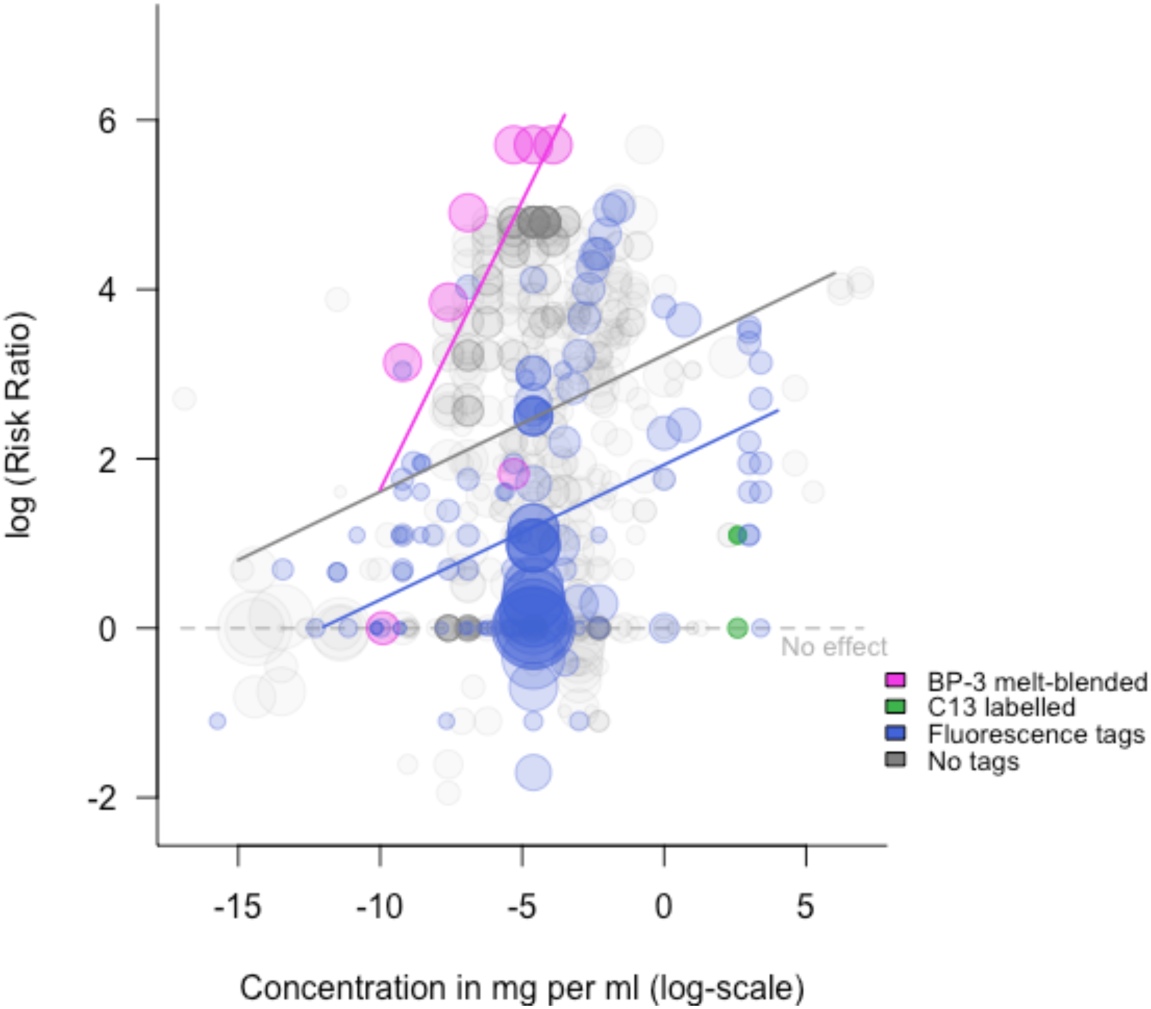
Log(Risk Ratios) of immobilization rates of daphnids exposed to NMP with and without additives or fluorescence tags. Point sizes reflect the log inverse standard errors of the data points. Meta-regression lines are shown for groups with at least five data points. BP-3: benzophenone-3.

#### 5.3.5 Uncontrolled surface modifications

Dissolved organic carbon (DOC) was added in five studies included in the meta-analysis, contributing 184 data points to the dataset. Three of these studies added humic acid (42 data points), one study added fulvic acid (18 data points) and two studies exposed daphnids in water sampled from natural lakes (124 data points). In addition, in two studies, NMP were incubated in medium containing dissolved organic carbon prior to using them in exposure bioassays. Incubation media were humic lake water, artificial medium containing humic acid, river water from the city centre of Ljubljana (Slovenia), spring water, water from close to a landfill site and water collected near a wastewater treatment facility (one data point each). The addition of DOC to the medium during exposure did not affect immobilization risk ratios consistently (Fig. 15). Compared to experiments without the addition of DOCs, risk ratios were slightly higher in experiments that exposed daphnids in natural lake water or added humic acid to the test medium. In contrast, they were, on average smaller when fulvic acid was added during exposure. Incubation of NMP in DOC-containing media prior to their use in exposure bioassays seems to have reduced the negative effects of NMP, but data is too limited at this point to draw clear conclusions.

**Figure 15.**
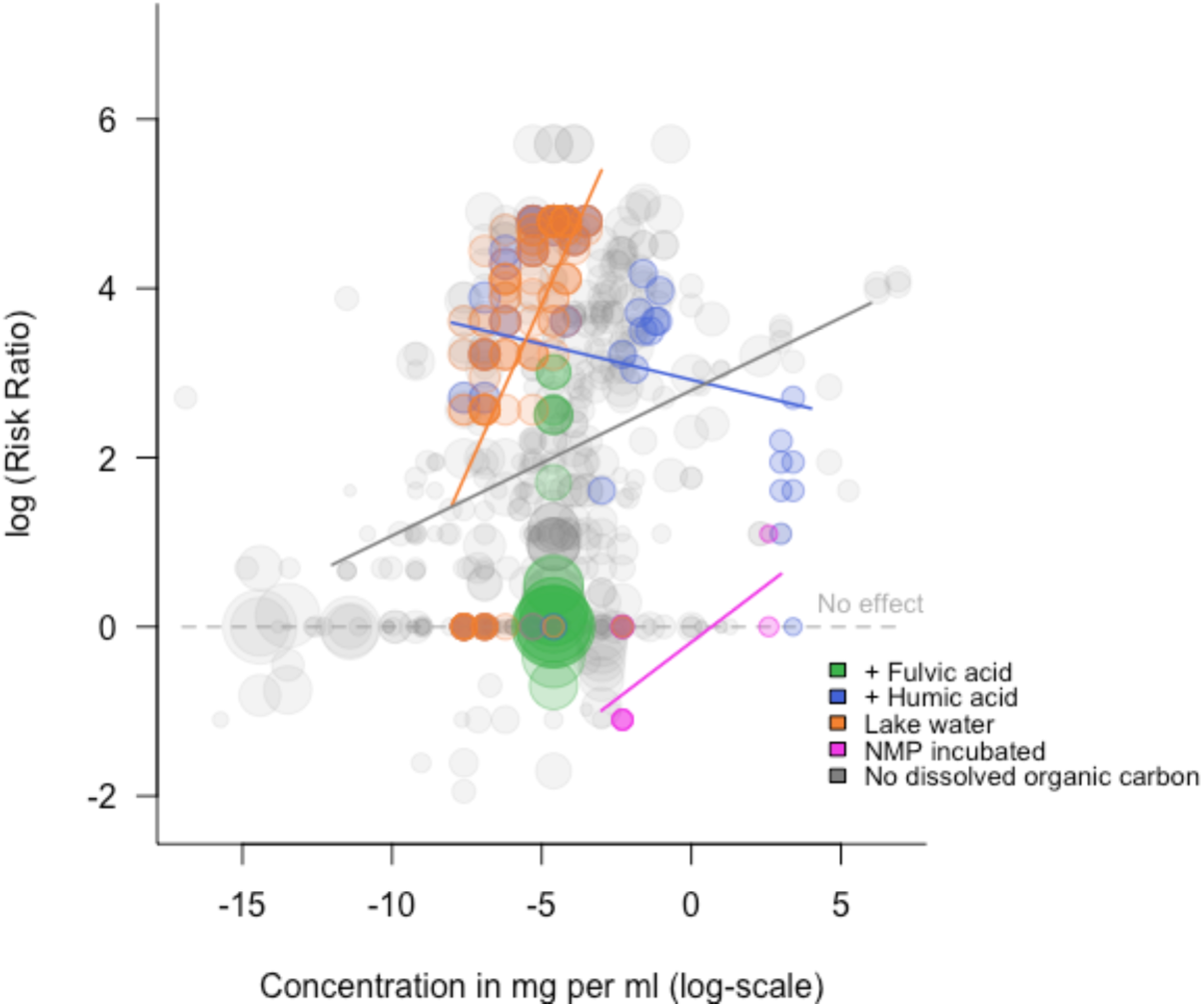
Log(Risk Ratios) of immobilization rates of daphnids exposed to NMP with and without the addition of dissolved organic carbon (DOC) in the medium during exposure or to NMP incubated in media containing DOC prior to exposure. Point sizes reflect the log inverse standard errors of the data points. Meta-regression lines are shown for groups with at least five data points.

#### 5.3.6 Controlled surface modifications

In our meta-analysis, eight studies reported the use of carboxylated NMP (63 data points), eight reported the use of aminated NMP (72 data points), one study reported the use of amidine derivatized NMP (190 data points) and three studies reported that they extracted the NMP prior to their use (15 data points).

Amidine-derivatized NMP seem to elicit a higher immobilization risk as compared to non-modified particles (Fig. 16). All data for amidine derivatized particles, however, have been extracted from only one study and validation studies would be needed to confirm (or disprove) whether the increased immobilization risk is indeed causally connected to the applied surface modification or resulted from other confounding parameters. The compiled data revealed no difference in immobilization risk ratios between aminated NMP, carboxylated NMP and NMP not modified prior to exposure. Surprisingly, NMPs extracted before being used in bioassays led to slightly higher immobilization risk ratios than unmodified NMPs.

**Figure 16.**
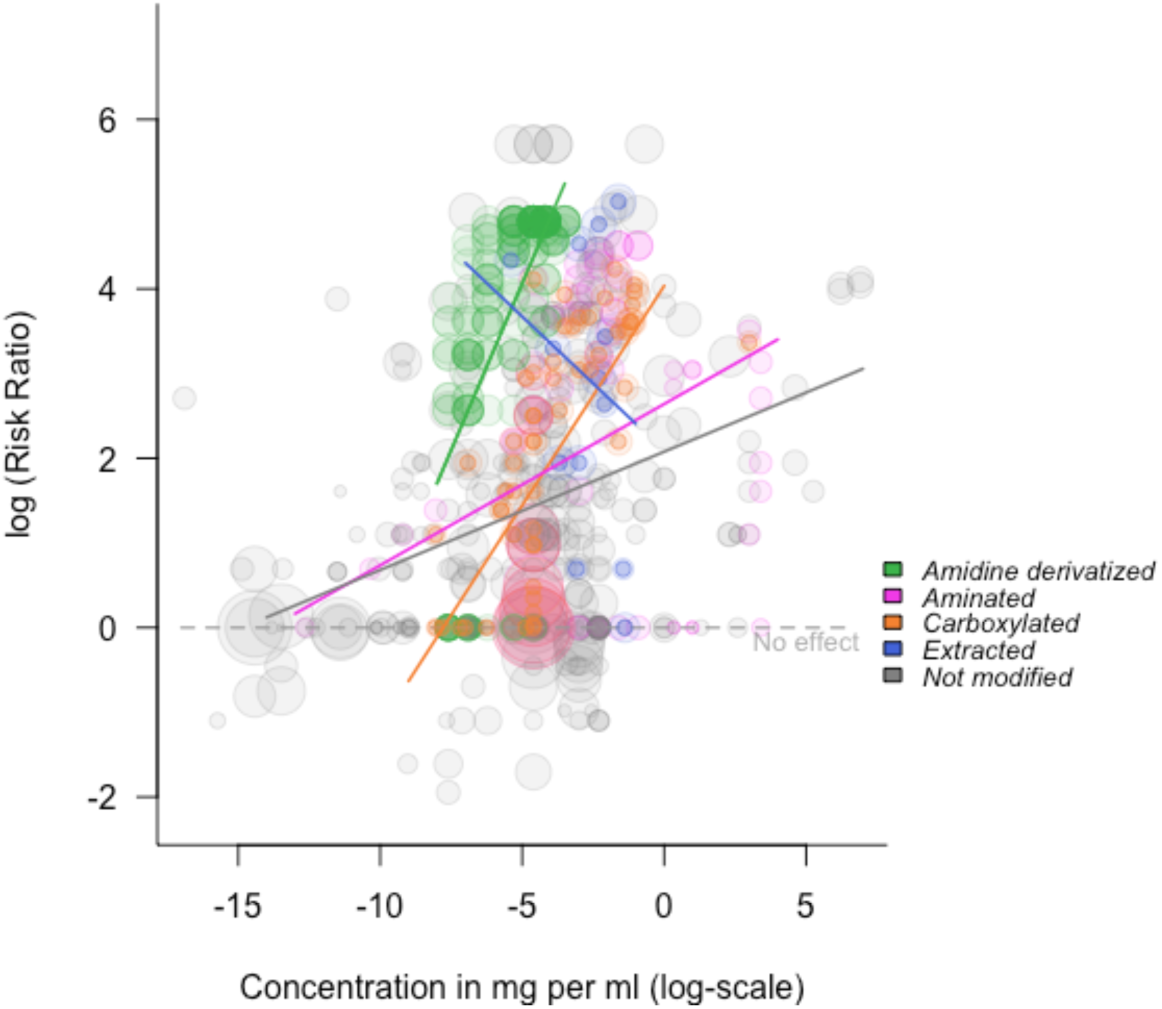
Log(Risk Ratios) of immobilization rates of daphnids exposed to NMP with different controlled surface modifications. Point sizes reflect the log inverse standard errors of the data points.

### 5.4 Prediction of NMP toxicity

NMP properties, experimental conditions and characteristics of the test organisms were sufficient to explain more than 90% of the variance in the data. In accordance with this, the predictive errors (residual mean squared errors, RMSE) for predictions on test datasets (from ten cross-validation splits) were consistently lower for the true data model than for the baseline model (Figure 17A). Predictor gains (relative contributions of predictors to the model) indicate that the concentration of NMP and the types of controlled modifications (i.e., controlled surface modifications like amination, carboxylation and amidine derivatization, and the addition of additives) may be important predictors for NMP toxicity (Figure 17 B). When looking at single feature gains (i.e., one feature per factor level; see Figure S6), we found that among NMP modification types, amidine derivatization seemed to be most important for achieving accurate predictions. Although controlled NMP modifications were most important for predictive accuracy, parameters related to the formation of a protein corona, ecocorona or biofilm (e.g., DOC added during exposure or incubation of particles in natural lake water) were used more frequently in splits (see predictor frequencies, Figure S7; for the predictors’ cover see Figure S8), indicating that these features were more often involved in fine tuning predictions. In addition to these parameters related to the surface structure of NMP, several other parameters seemed to be associated with NMP toxicity in the data, including NMP properties (e.g., NMP size, polymer type and density), experimental parameters (e.g., exposure duration) and properties of the test organisms (e.g., age).

**Figure 17.**
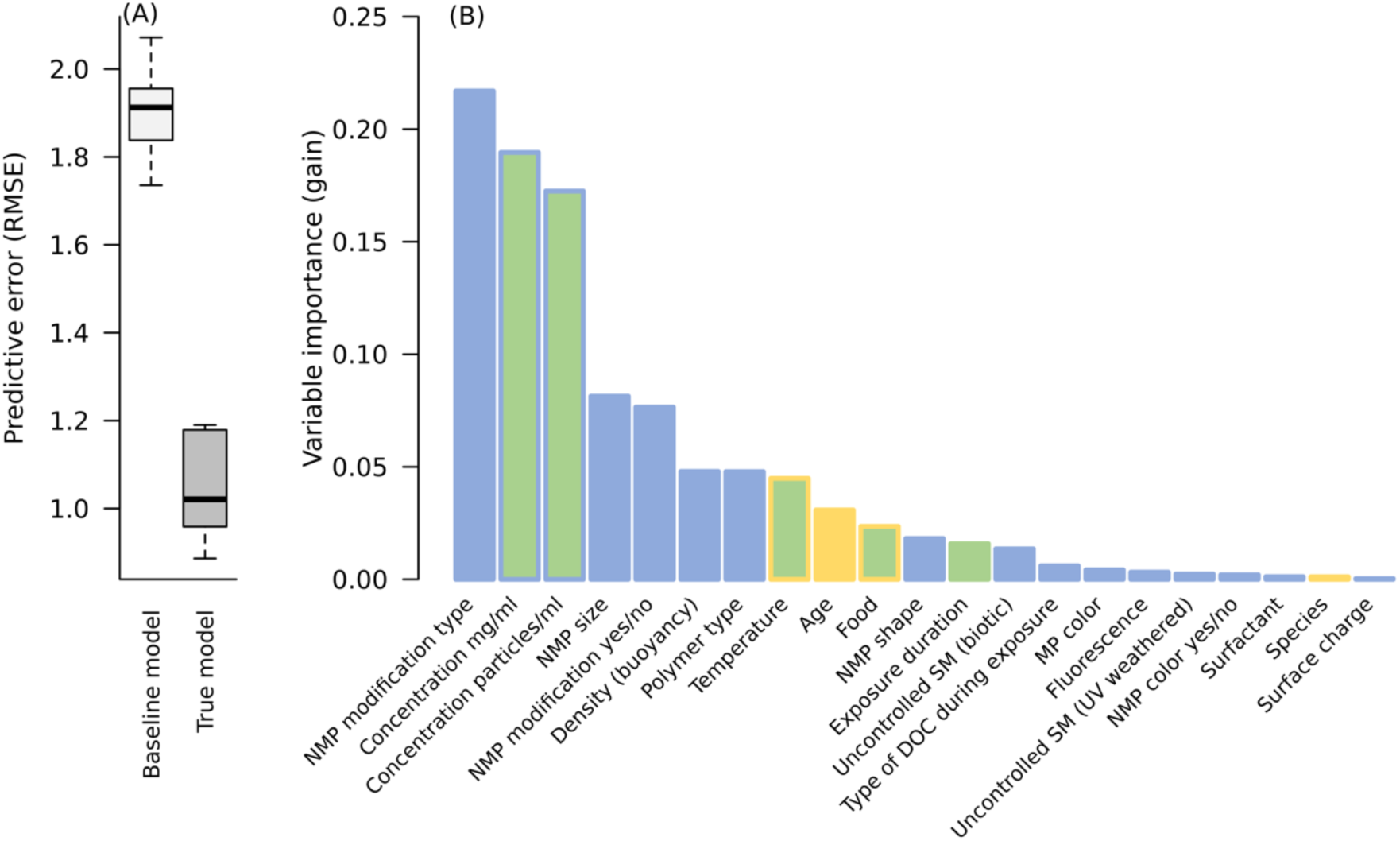
(A) Residual mean squared errors (RMSE) of boosted regression tree models predicting log Risk Ratios for immobilization for different NMPs based on NMP properties, experimental conditions and characteristics of the test organisms (n = 10 cross-validation splits). Baseline model: Model trained on data with randomly shuffled response values; True model: Model trained on true data. (B) Feature importances (gain) for predictor variables from the final boosted regression tree model trained on the full dataset. The colours in B correspond to the mindmap (Fig. 1).

## 6. Conclusions and Recommendations

This review article including a meta-analysis on immobilization aimed to provide an overview of which factors may influence test results of experiments with *Daphnia* and NMPs. We further depict the main NMP properties likely associated with adverse effects on *Daphnia*. Although we did not weigh the various NMP studies in terms of their quality, which was not the focus of our review, we want to note some key aspects here. In general, there is a consensus in the literature that methods in NMP research must be improved and harmonized, and several review articles have already addressed these challenges [e.g., 98,172–175]. Quality criteria identified include particle characteristics (i.e. particle size, shape, and polymer type; experimental design (i.e., chemical purity, laboratory preparation, verification of exposure); applicability in ecological risk assessment (i.e., endpoints, presence of dietary (natural) particles, effect thresholds); and ecological relevance (i.e., concentration, exposure time, ageing)[107].

Further, we would also like to emphasize that as much information as possible should be given about the organism and test conditions used. Only if these quality criteria are met, and all relevant information is given comparable and reliable data on the effects of NMP on *Daphnia* can be obtained. In addition, more attention should be paid to the biology of the test organisms. *Daphnia* are suitable and sensitive model organisms. However, species and clonal differences as well as their applicability for testing particulate contaminants, given the occurrence of a peritrophic membrane, should be considered or discussed with care, e.g. when studying tissue translocation of particulate contaminants.

In addition to these study quality criteria, the multidimensionality of NMP particles [208] is often not reflected in *Daphnia* research. Research gaps include using environmentally relevant exposure concentrations, using polymer types other than PS, shapes other than spherical shapes, and a combination of distinct NMP properties. As it becomes increasingly apparent that other NMP properties besides polymer type, shape, and size are additionally responsible for eliciting adverse effects, a more comprehensive characterization of particle properties is needed for hazard assessment. Here, especially surface charge, residual monomer content, additives and surface modifications should be considered and reported. A further mandatory prerequisite for particle toxicity testing should be the application of reference particles. We recommend using natural control particles as identical as possible in shape, size and density to the NMPs used. Interdisciplinary collaboration is needed to overcome the difficulties in adequately characterizing NMPs and obtaining NMPs with unique trait combinations. Ultimately, similar to the use of soluble reference substances such as potassium dichromate (K_2_Cr_2_O_7_) in ecotoxicity testing of chemicals (e.g. ISO standard 6341)[106], comparability and quality of NMP studies across laboratories could be improved by establishing standardized NMP reference particles that can be tested repeatedly in all experiments as a measure of quality control. Finally, there is a general lack of studies comparing the effects of specific single NMP traits, i.e. experiments where all properties except one are kept constant.

In summary, the multidimensionality that needs to be addressed in NMP research with *Daphnia* is currently not met in experiments. The literature review shows a scarcity of species-related diversity and NMP-related diversity, making it difficult to draw meaningful, environmentally relevant conclusions. The lack of species diversity could be explained by the fact that *D. magna* is the primary test organism in standard toxicity test guidelines, while the lack of diversity of polymer types and shapes may be explained by the lack of access to environmentally relevant particles. The lack of environmental relevance is also true for the concentrations used, commonly not reflecting reported environmental concentrations.

Further, our review revealed that there was a lack of relevant information, ranging from the organism of study to the setup of the experiment/exposure, and the NMPs used. Therefore, reaching a consensus on annotations and information that should be implemented in method sections that allow reproducing experiments is of great significance. Therefore, we her suggest a data sheet on the minimum reporting criteria when testing NMP effects on *Daphnia* (Table S2).

Nevertheless, thanks to the tremendous amount of reliable experimental data, we were able to draw first meaningful conclusions on which NMP properties may be the drivers of immobilization in *Daphnia*. Based on the data available, our meta-analysis indicates already two (three) NMP properties, concentration and polymer type (and density), plus the duration of exposure, to be the most relevant parameters for NMP toxicity in *Daphnia*. However, if research on NMP includes more and more detailed property combinations, not only could a better hazard assessment be made, but it would also provide a starting point for the future on how plastic products, that have a high probability of becoming MP, should be modified.

## Acknowledgements

We want to thank Rana Al-Jaibachi, Amanda Callaghan, Nathalie Tufenkji, Elvis Genbo Xu, Zhiquan Liu, Vera Slaveykova and Juan Saavedra for kindly providing raw data to their studies for the immobilization meta-analysis. We further thank Maysan Nashashibi for the photographs of the three daphnid species, and Robert Sigl for providing the photograph of the grazing *Daphnia*.

## Data availability

All data and code pertaining to this manuscript are available via Zenodo (XXX) and github (XXX). All data is also available in the online supplementary material at XXX.

## Author contribution

J.B., S.R., C.L. and M.M. designed the study. J.B. and S.R. performed the systematic review. J.B., S.R. and M.M. extracted immobilization data for the meta-analysis. J.B and M.M. performed data analysis and visualization. C.L. provided instruments and expert advice. All authors wrote, read and commented on the manuscript.

---

**Figure S1.**
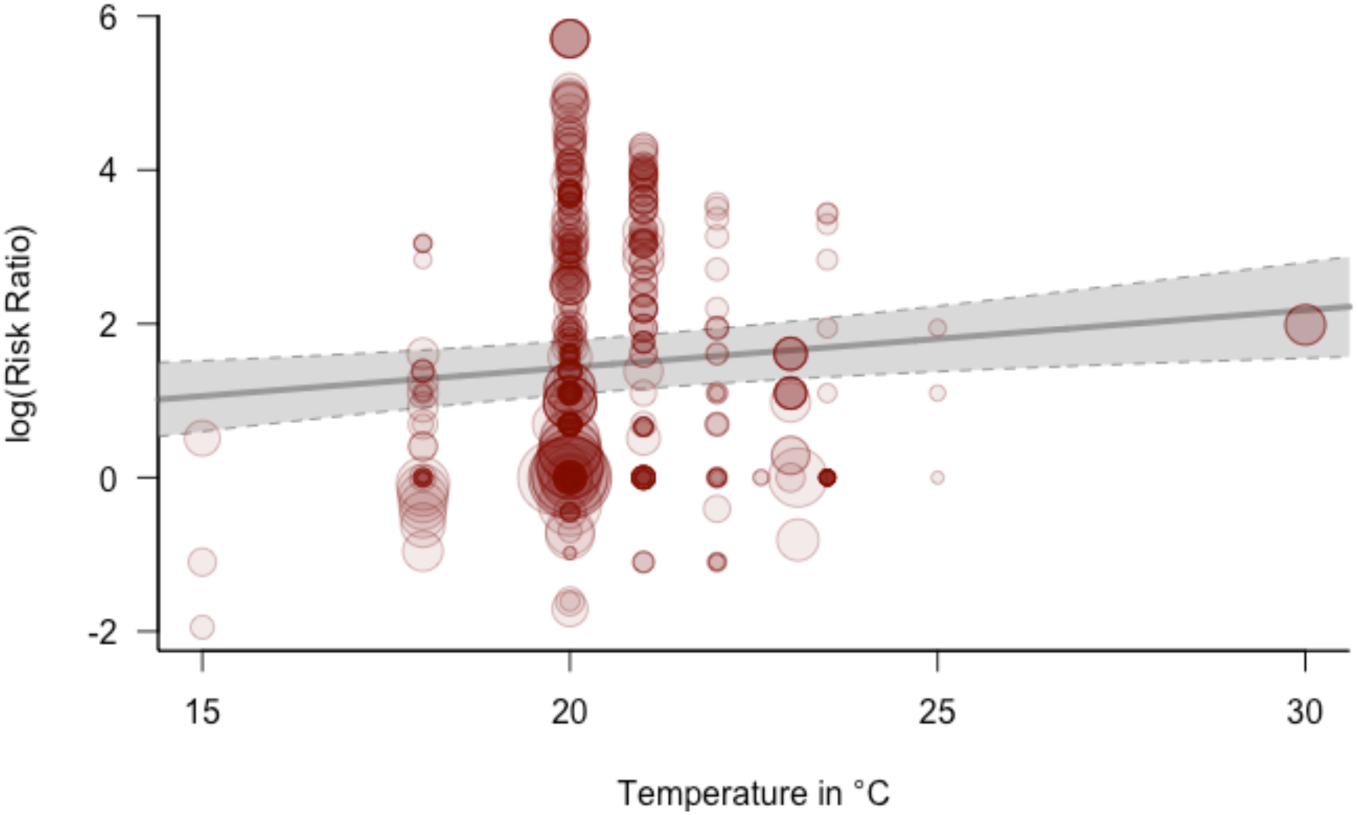
PRISMA flow diagram displaying the search results and the results from the publication screening process.

**Figure S2.**
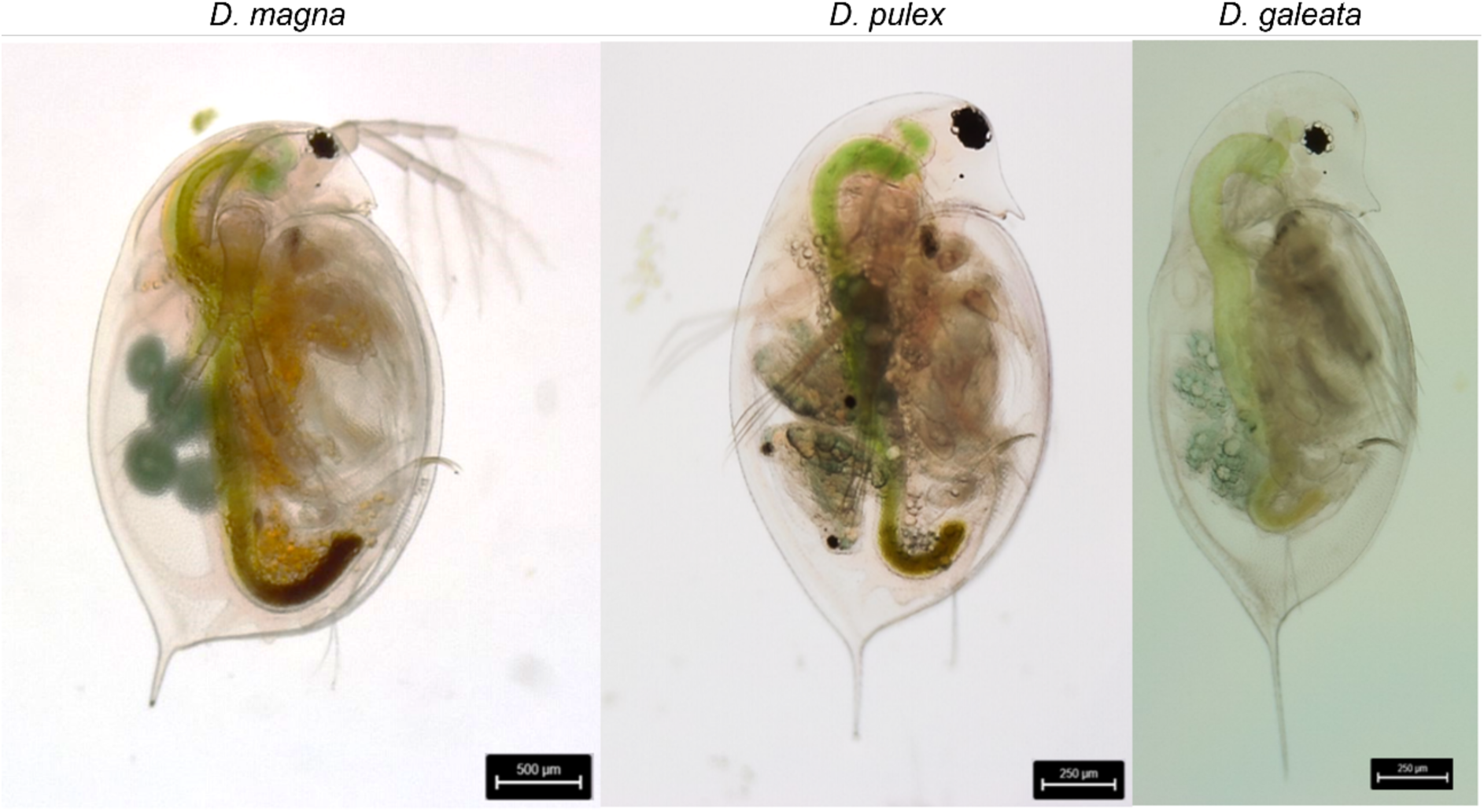
Bright-field microscopic images of adult representatives of the species *D. magna*, *D. pulex* and *D. galeata*, which are the species predominantly used in NMP research.

**Figure S3.**
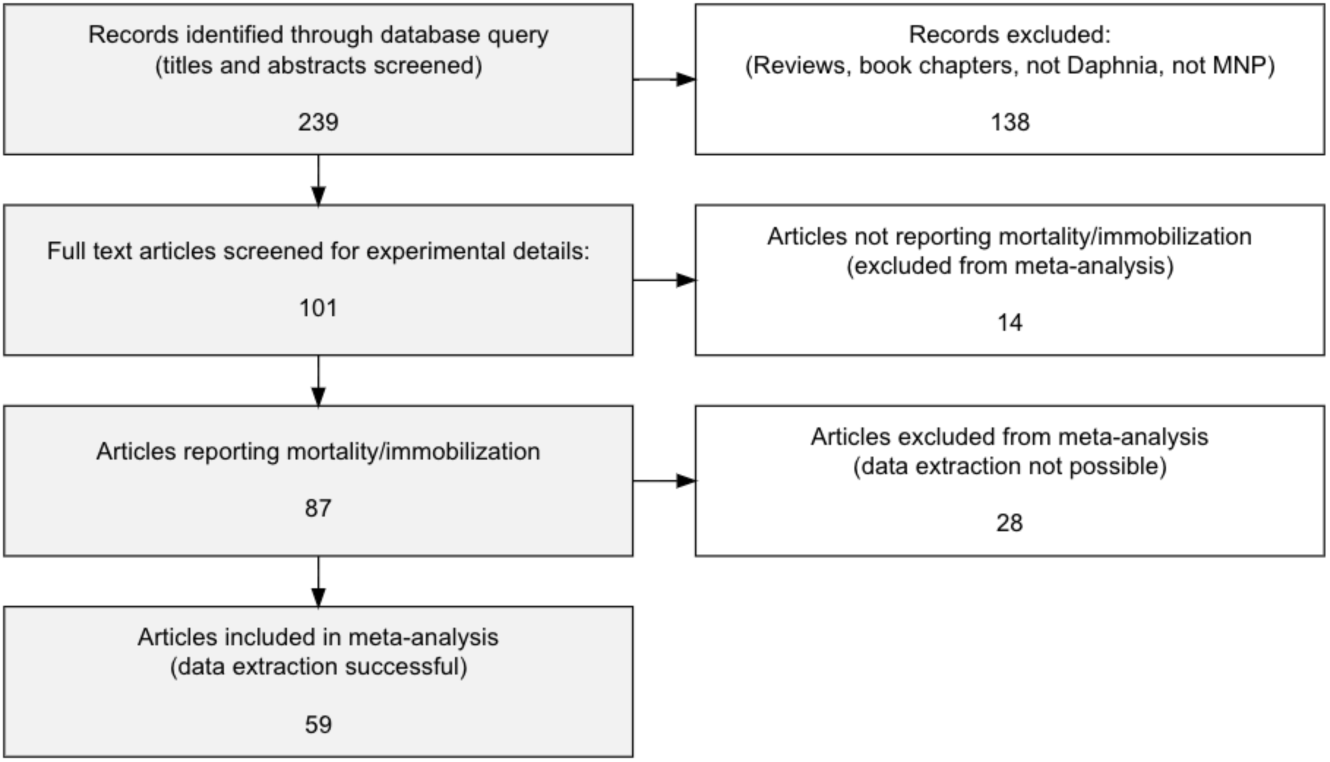
*D. magna* grazing on submerged macrophyte.

**Figure S4.**
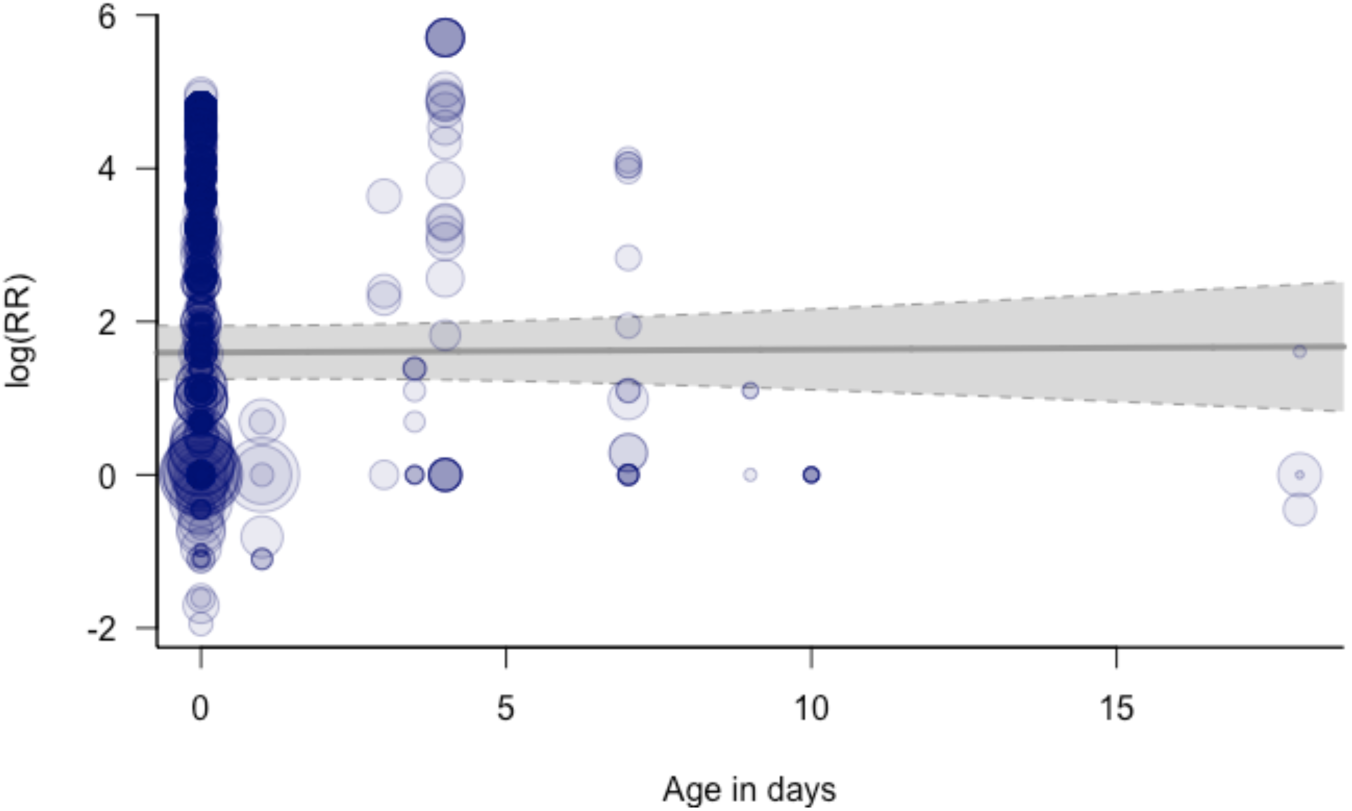
Influence of the age of test organisms at the start of exposure on the log(RR) for immobilization in *Daphnia* spp. exposed to NMP. Point sizes reflect inverse standard errors of the data points. The line represents the meta-regression line, and the shaded area represents the 95% confidence interval.

**Fig. S5:**
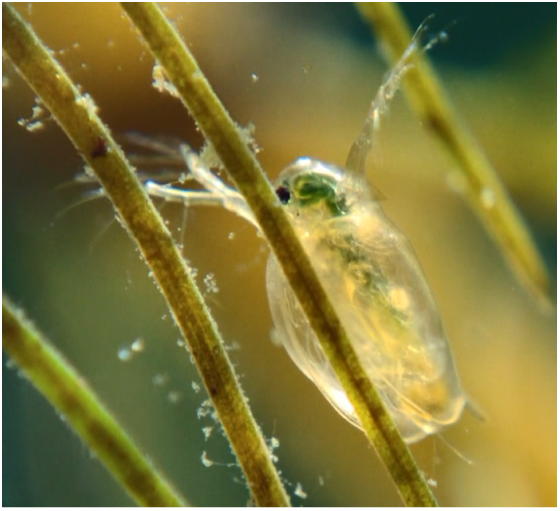
Influence of experimental temperature on log(Risk Ratio) of immobilization in *Daphnia* spp. exposed to different NMPs. Point sizes reflect inverse standard errors of the data points. The line represents the meta-regression line, and the shaded area represents the 95% confidence interval.

**TableS1:** Polymer types included in the meta-analysis of immobilization risk ratios in Daphnia spp. For each polymer type, the number of studies and the number of extracted data points are presented.

| Polymer name | Abbreviation | Number of studies | Number of data points |
| --- | --- | --- | --- |
| Low-density polyethylene | LDPE | 1 | 6 |
| Polyethylene | PE | 12 | 68 |
| Polyethylene terephthalate | PET | 4 | 11 |
| Mix of polyethylene, polystyrene and acrylonitrile butadiene styrene | PET/PS/ABS | 1 | 4 |
| Polyhydroxybutyrate | PHB | 1 | 8 |
| Poly(lactic acid) | PLA | 1 | 6 |
| Mix of polystyrene and latex | PS/Latex | 1 | 34 |
| Poly(methyl methacrylate) | PMMA | 1 | 8 |
| Poly(methyl methacrylate-co-stearyl methacrylate) copolymer | PMMA-PSMA | 1 | 8 |
| Polypropylene | PP | 2 | 5 |
| Polystyrene | PS | 31 | 488 |
| Polyurethane | PUR | 1 | 6 |
| Polyvinylchloride | PVC | 2 | 10 |
| Mix of polyvinylchloride and polyethylene | PVC/PE | 1 | 4 |
| One of ethylene acrylic acid, polyamide, a mix of polyethylene terephthalate and polyamide or undefined | other (one of EAA, PA, PET/PA or undefined) | 8 | 24 |

**Table S2:** Minimum information requirements for experiments with Daphnia and NMP, to be able to draw meaningful conclusions and to increase reproducibility of experiments.

|  |  |
| --- | --- |
| Daphnia Biology / Organismic level | Species<br>Clone / Sensitivity to reference chemicals<br>Age (start of experiment + when measuring endpoints)<br>Sex<br>Brood number |
| NMP level | Polymer type (+ manufacturer/preparation method)<br>Concentration (as particle count and mass count)<br>Shape<br>Size<br>Information on used control particles<br>Incubation method<br>Information on additives<br>Information on surface modifications (amination, carboxylation, Tween etc.) |
| Experimental level | Information on treatments<br>Duration<br>Temperature<br>Food quality and quantity<br>Replicates<br>Animals/Replicate<br>Information on used media (also quantities) |

